# Hierarchical Coordination of Polymerase Theta and RAD51 Resolves Clustered Replication Fork Collapse

**DOI:** 10.1101/2025.07.02.662846

**Authors:** Chelsea M. Smith, Dennis A. Simpson, Paolo Guerra, Wanjuan Feng, Simon W. Ellington, Dongbo Lu, Carel Fijen, Natalie M. Boylston, Kishore K Chiruvella, Iman El-Shiekh, Rachel Lee, Anne-Sophie Wozny, Andrew M. Pregnall, Brian E. Eckenroth, Afaf H. El-Sagheer, Tom Brown, Sylvie Doublié, Dale A. Ramsden, Eli Rothenberg, Serena Nik-Zainal, Gaorav P. Gupta

## Abstract

The essential role of polymerase theta (Polθ)-mediated end joining (TMEJ), an alternative double strand break repair pathway, has been primarily studied in homologous recombination (HR)-deficient contexts(*1*, *2*). Here, we uncover an indispensable role for TMEJ in HR-proficient mammalian cells during the repair of interstrand crosslinks (ICLs). We show that Polθ is recruited downstream of canonical ICL repair steps—including ICL unhooking, RAD51 loading, and RAD51 ubiquitylation—and localizes to sites of unresolved HR through interactions with ubiquitylated RAD51 filaments. Using genomic scar profiling and targeted ICL repair assays, we find that TMEJ resolves a minor subset of lesions that are not amenable to HR repair, such as clustered ICLs that can induce two-ended replication fork collapse. These findings reveal a RAD51 ubiquitylation-dependent mechanism for Polθ recruitment and establish TMEJ as a hierarchically deployed DNA repair pathway that safeguards genome stability when HR is insufficient to resolve replication-associated DNA damage.

**Short Summary:** Polθ is recruited via RAD51 ubiquitylation to resolve clustered ICLs that generate HR-refractory replication fork collapse.

## Background

DNA replication-associated double strand breaks (DSBs), arising from replication fork collapse, represent a major threat to genome integrity(*3*). Homologous recombination (HR) is the dominant and most accurate mechanism for repairing replication-associated DSBs(*4*). Recently, polymerase theta (Polθ) mediated end joining (TMEJ) has emerged as an alternative, inherently mutagenic pathway that also repairs replication-associated DSBs(*5*, *6*). During TMEJ, Polθ scans resected 3’ single-stranded DNA (ssDNA) overhangs for short stretches of complementary sequence (i.e., microhomology), conducts nucleolytic processing and templated DNA synthesis in coordination with DNA polymerase delta, and culminates in ligation of the broken DNA ends(*7*). In contrast, HR requires more extensive end resection to produce long (>100 nucleotides) 3’ ssDNA overhangs that are loaded with RAD51 recombinase, enabling use of a homologous sister chromatid template for error-free repair(*8*). Because TMEJ introduces microhomology-associated deletions and/or templated insertions, its activity must be restricted to a minority of DSB repair events. However, how Polθ activity is suppressed to favor HR, despite TMEJ requiring less DSB end resection, remains poorly understood.

The essential role of Polθ/TMEJ has predominantly been characterized in cells deficient for HR, non-homologous end joining (NHEJ), or other canonical repair pathways(*1*, *2*, *9*, *10*). In these contexts, TMEJ is considered a backup repair mechanism, implicated in the repair of daughter strand gaps, two-ended DSBs, and replication-associated DSBs that persist into mitosis(*9*, *11–14*). This dependency of HR-deficient cancers on TMEJ underlies the pursuit of Polθ inhibitors for synthetic lethality-based therapy(*15*).

Recent studies have begun to elucidate the regulation of Polθ, though significant knowledge gaps remain. While Polθ/TMEJ exhibits activity in S phase(*13*, *16*, *17*), Polθ has also been implicated in mitotic DSB repair, with RHINO-dependent recruitment during mitosis and Plk1-mediated phosphorylation promoting Polθ-TOPBP1 interactions(*12*, *18*). However, these studies largely focus on HR-deficient contexts and do not resolve how HR and TMEJ are hierarchically coordinated under HR proficiency.

The evolutionary conservation of Polθ across plants and metazoans suggests that Polθ fulfills essential roles even in HR-proficient cells(*11*, *19*). Polθ-deficient cells exhibit spontaneous DNA damage and heightened sensitivity to replication stress-inducing agents(*10*, *20*), arguing that TMEJ does not merely compete with HR but is uniquely required to resolve specific types of replication-associated DNA damage. In *C. elegans*, TMEJ has been shown to protect collapsed replication forks from extensive deletions and rearrangements(*21*). However, replication-associated DSBs are an optimal substrate for HR, and the distinct profile of replication-associated lesions that necessitate TMEJ remains undefined.

DNA interstrand crosslinks (ICLs) are among the most formidable barriers to replication fork progression, requiring the coordinated activity of the Fanconi Anemia (FA) pathway, homologous recombination (HR), nucleotide excision repair, and translesion synthesis (TLS) for resolution (Figure 1A)(*22–24*). Although Polθ has an evolutionarily conserved role in ICL repair(*10*, *25–29*), its molecular function within this process, particularly in HR-proficient cells, remains poorly understood. This gap is especially notable given Polθ’s dual identity as both a therapeutic target and a genome maintenance factor. Here, we leverage Mitomycin C (MMC)-induced ICLs as a tool to define the regulatory context, recruitment mechanism, and functional contribution of Polθ during replication-associated DNA repair.

**Figure 1.**
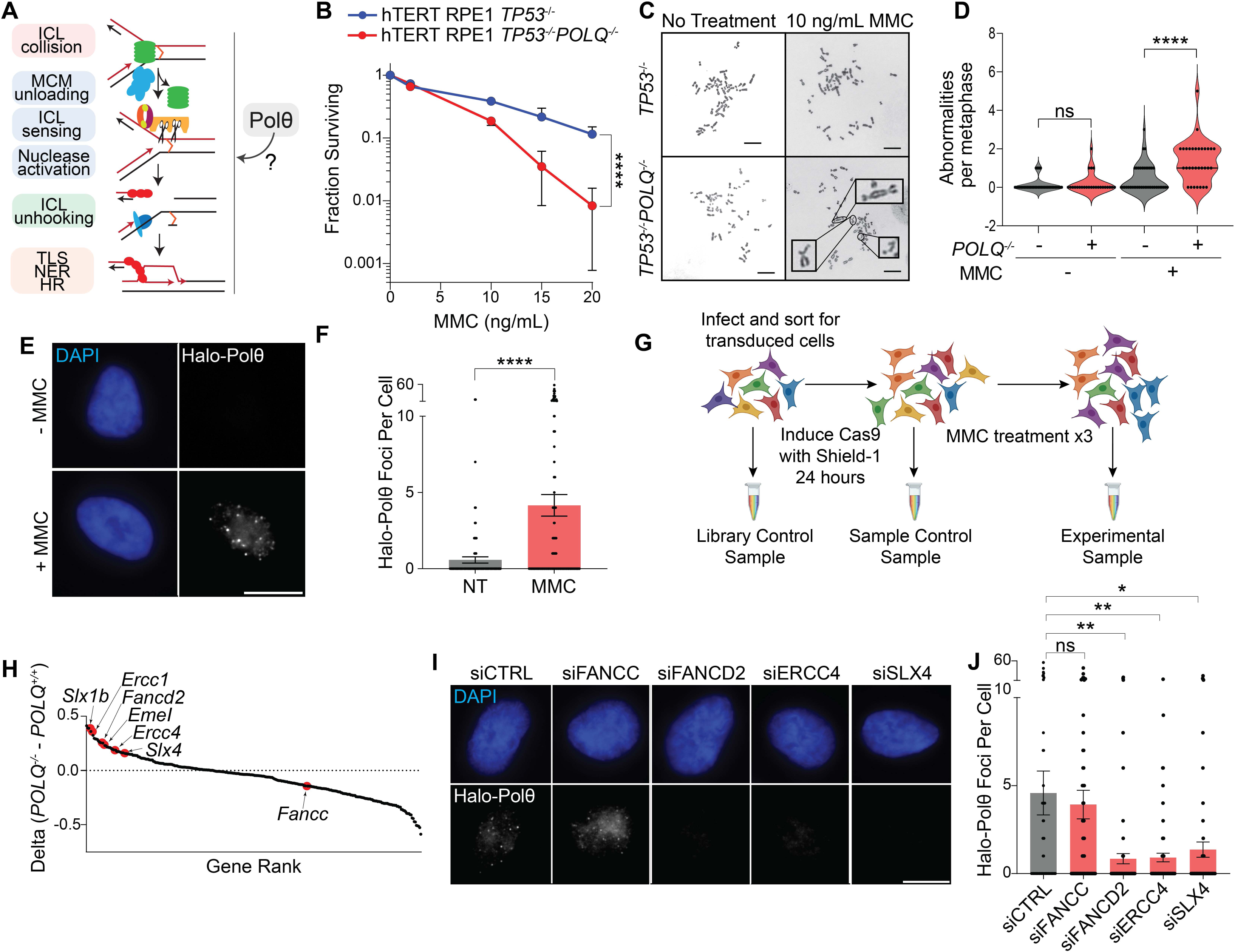
Role of Polθ in ICL repair and genome maintenance. (A) Schematic overview of interstrand crosslink (ICL) repair pathways and possible roles of Polθ. (B) Colony formation assay demonstrating increased mitomycin C (MMC) sensitivity in hTERT RPE1 *TP53^− /-^POLQ^−/-^* cells relative to *TP53^−/-^* controls. Statistical significance: ****P < 0.0001. (C) Representative metaphase spreads of hTERT RPE1 *TP53^−/-^* and *TP53^−/-^POLQ^−/-^* cells treated with 10 ng/mL MMC for 15 h. Insets highlight chromatid breaks, telomere associations, and dicentric chromosomes. (D) Quantification of chromosomal aberrations in *TP53^−/-^* and *TP53^−/-^POLQ^−/-^* cells after MMC treatment. ****P < 0.0001, unpaired t-test. (E) Representative images of Halo-tagged Polθ in hTERT RPE1 cells labeled with Halo-Janelia Fluor 646, ±100 ng/mL MMC for 6 h. Images show DAPI (blue) and Halo signal (grey). (F) Quantification of Halo–Polθ nuclear foci per cell. MMC treatment significantly increased Polθ foci (****P < 0.0001, Welch’s t-test). (G) Schematic of the CRISPR screen design to identify genes epistatic with Polθ following three sequential 20 ng/mL MMC treatments. (H) Delta plots of sgRNA abundance change score (ACS) in Polq^−/-^ MEFs minus sgRNA ACS in Polq^+/+^ (i.e., Polq^−/-+hPOLQ^) MEFs for all 309 genes in the DDR-CRISPR library. Genes with positive values are epistatic with *POLQ* in MMC survival, and are enriched in factors required for ICL nuclease activity and recruitment. (I) Representative images of Halo–Polθ localization in cells treated with siRNAs targeting FANCC, FANCD2, SLX4, or ERCC4, ±100 ng/mL MMC (6h). DAPI (blue), Halo-Polθ (grey). (J) Quantification of Halo-Polθ foci per cell. MMC-induced Polθ foci were significantly reduced upon FANCD2 (<u>**P < 0.01</u>), ERCC4 (**P < 0.01), or SLX4 (*P = 0.02) knockdown; no significant change for FANCC (P = 0.66). Statistical comparisons by two-tailed Welch’s t-test. Data are shown as mean ± SEM. Scale bars, 10 µm.

## Results

### Polymerase Theta Protects Cells from MMC Induced Death

Polε has been implicated in ICL repair in multiple organisms(*10*, *25–27*). To examine the role of Polθ in human cells, we generated *TP53^−/-^ POLQ^−/-^* hTERT-RPE1 cells and assessed their sensitivity to MMC using a clonogenic survival assay. *POLQ*-deficient cells exhibited significantly increased sensitivity to MMC compared to parental *TP53^−/-^*controls (Figure 1B).

In contrast to cells deficient for components of the Fanconi anemia (FA) pathway, which form Polθ - dependent radial chromosomes upon MMC exposure(*30*), *POLQ^−/-^* cells displayed increased chromatid breaks and fusions without radial formation, consistent with prior observations in *Polq^−/-^* MEFs (Figure 1C-D)(*10*). To visualize Polθ localization during ICL repair, we engineered hTERT-RPE1 cells to constitutively express N-terminal Halo-tagged Polθ (hTERT-RPE1^Halo-Polθ^) and confirmed expression by Western blotting and RT-PCR (Supplementary Figure 1A-1B). Following MMC treatment, Halo-Polθ formed discrete nuclear foci (Figure 1E-F), indicating recruitment to sites of DNA damage. Additionally, MMC-induced Halo-Polθ foci were predominantly formed during both early and late S phase, with no appreciable increase in M phase cells (Supplementary Figure 2A-D). These findings suggest that Polθ recruitment in S phase promotes survival following ICL induction through a mechanism that is at least partially distinct from canonical FA pathway repair.

### Polθ is Epistatic with Genes Required for ICL Unhooking

To define the functional relationship of Polθ within the ICL repair pathway, we conducted a CRISPR-based genetic interaction screen. *Polq^−/-^*MEFs, with or without reconstitution with human *POLQ*, were transduced with a CRISPR library targeting 309 DNA repair genes and subjected to sequential MMC treatments (Figure 1G). Small guide RNA (sgRNA) depletion analysis using the Völundr pipeline identified multiple genes whose loss affected cell survival in both Polθ-deficient and reconstituted contexts (Supplementary Figure 1C)(*10*). By plotting the difference in sgRNA depletion scores between the absence or presence of Polθ, we identified genes that are epistatic to Polθ in MMC survival, indicating that the gene’s contribution to survival is at least partially Polθ-dependent (Figure 1H). Notably, the FA pathway genes specifically required for ICL unhooking - *Fancd2*, *Eme1*, *Ercc1*, *Slx1b*, *Ercc4*, and *Slx4* – were all identified as epistatic with Polθ. In contrast, many components of the FA core complex that are not specifically implicated in ICL unhooking, such as *Fancc*, did not exhibit epistasis with Polθ.

To validate these epistatic relationships, we depleted *FANCC*, *FANCD2*, *ERCC4*, and *SLX4* individually in hTERT-RPE1^Halo-Polθ^ cells using siRNA. Knockdown efficiency was confirmed by RT-PCR (Supplementary Figure 1D-G). Following MMC treatment, immunofluorescence analysis revealed that Halo-Polθ foci formation was markedly reduced upon FANCD2, ERCC4, or SLX4 depletion, but not after FANCC knockdown (Figure 1I-J). These findings build on prior work demonstrating that FANCD2 is required for Polθ/TMEJ engagement at stalled replication forks in BRCA-deficient settings(*31*), and extend this mechanism to HR-proficient cells undergoing ICL repair. Together, our findings position Polθ recruitment downstream of FANCD2-dependent nuclease activity, consistent with a role for Polθ in the repair of replication-associated DSBs generated after ICL unhooking.

### Polθ Colocalizes with a Subset of RAD51 Foci after MMC Induced DNA Damage

To investigate the spatial relationship between Polθ and RAD51 during DNA damage repair, we analyzed Halo-tagged Polθ colocalization with RAD51 following MMC treatment (Figure 2A). Under basal conditions (DMSO), colocalization was minimal (Figure 2B). In contrast, MMC treatment substantially increased colocalization, with ∼80% of Polθ foci overlapping RAD51 and ∼30% of RAD51 foci overlapping Polθ (Figure 2A-B). A similar pattern was observed following ionizing radiation (Supplementary Figure 3A-B). Super-resolution imaging of replication fork collapse induced by Top1 inhibition also demonstrated significant colocalization between Polθ and RAD51 (Supplementary Figure 3C-E). Quantitative analysis revealed that RAD51 foci colocalized with Halo-Polθ exhibited greater fluorescence intensity than non-colocalized RAD51 foci (Figure 2D), suggesting preferential recruitment of Polθ to more mature RAD51 nucleofilaments. These findings support a model wherein Polθ is selectively recruited to a subset of mature RAD51 nucleofilaments that persist following ICL unhooking and replication fork collapse.

**Figure 2.**
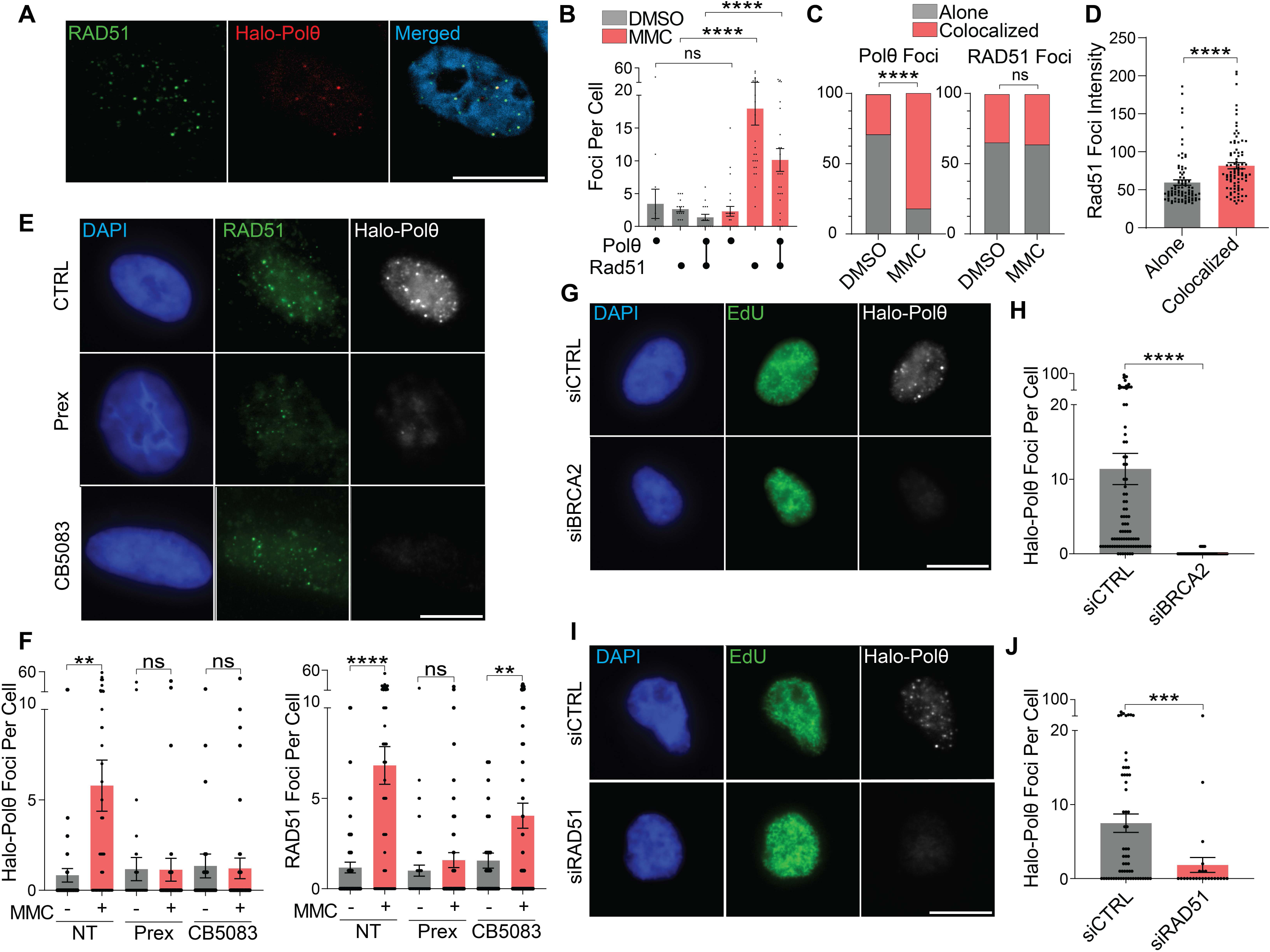
Hierarchical recruitment of RAD51 and Polθ following ICL damage. (A) Representative confocal images of Halo-Polθ and RAD51 in hTERT-RPE1^Halo-Polθ^ cells ±100 ng/mL MMC for 6h. RAD51 (green), HaloTag-JFX646 (red), and DAPI (blue). (B) Quantification of foci per nucleus. Colocalized RAD51-Polθ foci are significantly increased with MMC (***<u>*</u>P < 0.0001); Polθ-only foci not significantly altered (P = 0.8164). (C) Proportion of total foci colocalized for Polθ (left) and RAD51 (right) following DMSO or MMC treatment. Only Polθ demonstrates a significant increase in colocalization (***P<0.0001 vs. P=0.89; Fisher’s exact test). (D) RAD51 intensity (arbitrary units) in single versus colocalized foci; colocalized foci showed significantly higher intensity (****P < 0.0001, Welch’s t-test). (E) Representative images after treatment with MMC ± p97 inhibitor CB5083 (1 µM) or Chk1 inhibitor Prexasertib (250 nM). DAPI (blue), RAD51 (green), Halo–Polθ (grey). (F) Quantification of Halo-Polθ (left) and RAD51 (right) foci per nucleus. CB5083 and Prexasertib significantly decreased foci compared to MMC alone (****P < 0.0001, **P < 0.01). (G) Representative images after siRNA knockdown of BRCA2. EdU (green), Halo–Polθ (grey), DAPI (blue). (H) Quantification of foci per EdU+ nucleus. BRCA2 knockdown significantly reduced Polθ foci (***<u>*</u>P < 0.00<u>0</u>1). (I–J) As in G–H, for RAD51 knockdown. RAD51 depletion significantly reduced Polθ foci per EdU+ nucleus (***P < 0.001). All data represented as mean ± SEM and use two-tailed tests. P values calculated using Welch’s t-test unless otherwise noted. Scale bars, 10 µm.

### MMC Induction of Polθ foci Requires CHK1 Signaling and CMG Helicase Unloading

To further evaluate the temporal relationship between RAD51 and Polθ recruitment, we examined early steps of replication fork processing after ICL induction, specifically the role of CHK1 activation and p97-dependent CMG helicase unloading(*23*). hTERT-RPE1^Halo-Polθ^ cells were treated with MMC in the presence or absence of CHK1 inhibitor (Prexasertib) or p97 inhibitor (CB-5083) for 6 hours followed by immunofluorescence for RAD51 and Halo-Polθ. Inhibition of CHK1, which orchestrates the replication stress response, impaired recruitment of both RAD51 and Polθ (Figure 2E-F). In contrast, inhibition of CMG helicase unloading through p97 blockade permitted at least partial RAD51 loading but significantly impaired Polθ recruitment (Figure 2E-F). These findings suggest a temporal hierarchy during ICL repair: RAD51 is loaded following CHK1-mediated replication stress signaling and, at least in part, prior to replication fork collapse, whereas Polθ recruitment predominantly occurs after CMG helicase unloading.

### Polθ Foci Formation is Dependent on RAD51 Loading

Our data suggested that RAD51 loading precedes Polθ recruitment during ICL repair. To directly assess whether RAD51 filament formation is required for subsequent Polθ recruitment, we depleted BRCA2—a critical mediator of RAD51 nucleofilament assembly at resected DSBs—using siRNA. Knockdown efficiency was confirmed by RT-PCR (Supplementary Figure 3F). Since BRCA2 depletion can impair cell cycle progression, we restricted our analysis to S phase cells to ensure observed effects were not secondary to cell cycle perturbations. Following a 6-hour MMC treatment, BRCA2 knockdown markedly impaired both RAD51 and Halo-Polθ foci formation (Figure 2G-H and Supplementary Figure 3G). RAD51 knockdown similarly reduced Halo-Polθ foci formation in S phase cells following MMC (Figure 2I-J). These findings indicate that RAD51 nucleofilament assembly is a critical substrate for Polθ recruitment in S phase following DNA damage-associated replication fork collapse.

### Ubiquitylation of RAD51 is Required for Polθ Recruitment

Because Polθ is recruited to only a subset of RAD51 foci, we hypothesized that a post-translational modification of RAD51 may trigger Polθ recruitment to sites of DNA damage. We focused on RAD51 ubiquitylation, which is induced at mature RAD51 nucleofilaments and has been implicated in regulating RAD51 unloading(*32*).

We initially targeted three E3 ligases known to ubiquitylate RAD51—RFWD3, FBH1, and FBXO5—with siRNA knockdown, and confirmed they were depleted by RT-PCR (Supplementary Figure 4A-D)(*32–34*). Knockdown of RFWD3 did not impair MMC-induced Halo-Polθ recruitment in S phase relative to siControl cells, whereas knockdown of FBXO5 or FBH1 significantly reduced Halo-Polθ foci formation. Combined knockdown of FBXO5 and FBH1 did not further diminish Polθ foci, suggesting functional redundancy and the potential involvement of additional factors in RAD51 ubiquitylation-dependent Polθ recruitment (Figure 3A-B).

**Figure 3.**
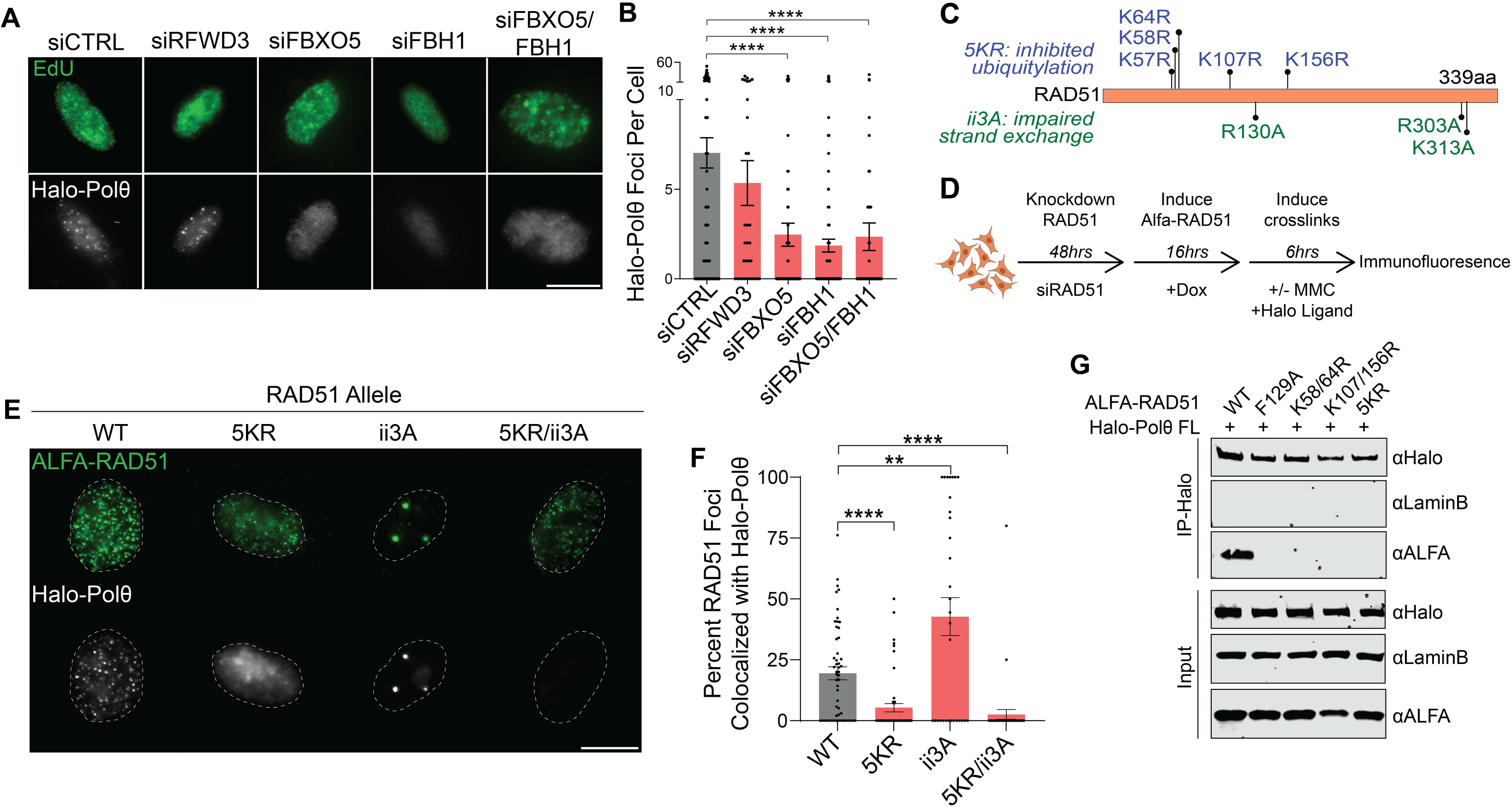
RAD51 ubiquitylation is required for Polθ recruitment after ICL damage. (A) Representative images of Halo-Polθ (grey) after siRNA knockdown of RFWD3, FBXO5, FBH1, or FBXO5+FBH1, after MMC (6h) in EdU (green) and siGLO positive nuclei.. (B) Quantification of Polθ foci per EdU/siGLO positive nucleus. FBXO5, FBH1, and dual knockdown significantly impaired Polθ recruitment relative to siCTRL (****P < 0.0001). (C) Schematic of RAD51 mutant alleles: 5KR (blue), ii3A (green). (D) Experimental strategy for inducible expression of siRNA-resistant RAD51 variants in hTERT-RPE1^Halo-Polθ^ cells with endogenous RAD51 knockdown. (E) Representative images of Halo-Polθ and ALFA-tagged RAD51 mutants after 100 ng/mL MMC (6 h). White hashed lines indicate the nuclear outline as defined by DAPI (not shown). (F) Quantification of colocalized foci in MMC treated cells. RAD51-5KR and 5KR/ii3A mutants reduced, while ii3A alone enhanced, Polθ– RAD51 colocalization (****P < 0.0001; **,P<0.01). (G) Nuclear extract co-immunoprecipitation of ALFA-RAD51 variants in HEK293T cells. Wild-type, but not RAD51 mutants, interact with Halo–Polθ. All data represented as mean ± SEM and used an unpaired, two-tailed Welch’s t-test.

To investigate the role of RAD51 ubiquitylation more directly, we generated hTERT-RPE1^Halo-Polθ^ cells expressing doxycycline (Dox)-inducible, siRNA-resistant ALFA-tagged RAD51 transgenes. These included RAD51-WT, RAD1-5KR (K57/58/64/107/156R, ubiquitylation deficient) ((*32*)51-ii3A (R130A/R303A/K313A, strand exchange deficient)(*35*), and a RAD51-5KR/ii3A compound mutant (Figure 3C). Following endogenous RAD51 knockdown and transgene induction, cells were treated with MMC and processed for immunofluorescence (Figure 3D). Expression of ALFA-RAD51 transgenes was confirmed by Western blotting (Supplementary Figure 4E). Expression of RAD51^WT^ restored RAD51-colocalized Halo-Polθ foci formation, whereas RAD51^5KR^ expression significantly reduced colocalization with Halo-Polθ. Expression of the HR defective allele RAD51^ii3a^ significantly increased Halo-Polθ colocalization, and the RAD51^5KR/ii3a^ compound mutant allele abolished this induction of RAD51-colocalized Halo-Polθ foci (Figure 3E-F). These findings indicate that Polθ recruitment to RAD51 requires ubiquitylation and is enhanced when HR is interrupted.

To further dissect the contribution of specific ubiquitylation sites and E3 ligases, we generated additional ALFA-tagged RAD51 mutants (K58/64R and K107/156R) that have been proposed as target lysines for RFWD3-, FBH1-, or FBXO5-dependent ubiquitylation, and a RAD51^F129A^ mutant that has been shown to disrupt interaction with FBXO5(*33*). To examine the effect of RAD51 ubiquitylation on its interaction with Halo-Polθ, HEK293T cells were co-transfected with Halo-Polθ and RAD51 mutant constructs, and co-immunoprecipitation (co-IP) analyses of nuclear extracts was performed. Whereas RAD51^WT^ efficiently co-immunoprecipitated with Halo-Polθ, none of the RAD51^KR^ mutants (5KR, K58/64R, K107/156R) nor RAD51^F129A^ interacted with Halo-Polθ (Figure 3G). Collectively, these findings highlight that RAD51 ubiquitylation at multiple sites, mediated by FBH1 and FBXO5, is essential for Polθ recruitment to sites of DNA damage-associated replication fork collapse.

### RAD51 and Ubiquitin Binding Domains in Polθ Required for RAD51 Association

To identify the specific regions of Polθ that mediate its interaction with RAD51, we initially examined the Polθ helicase-like domain, disordered central domain, and polymerase domain. We transiently co-overexpressed ALFA-RAD51 and Halo-Polθ constructs—including full-length wild-type (FL), helicase-like domain only (HLD), central domain deletion (ΔCEN), and nuclear localization signal-containing polymerase domain only (PolD)—in HEK293T cells (Figure 4A). Co-IP from nuclear extracts demonstrated that while the PolD construct retained RAD51 interaction, neither the HLD alone nor the ∆CEN construct showed detectable binding. Given that ∆CEN retains both HLD and PolD, its lack of interaction is unexpected and may reflect altered PolD accessibility when tethered to HLD in the absence of the central domain. These findings suggest that multiple Polθ domains contribute to RAD51 interaction and that central domain integrity may facilitate proper domain orientation or cooperativity required for binding.

**Figure 4.**
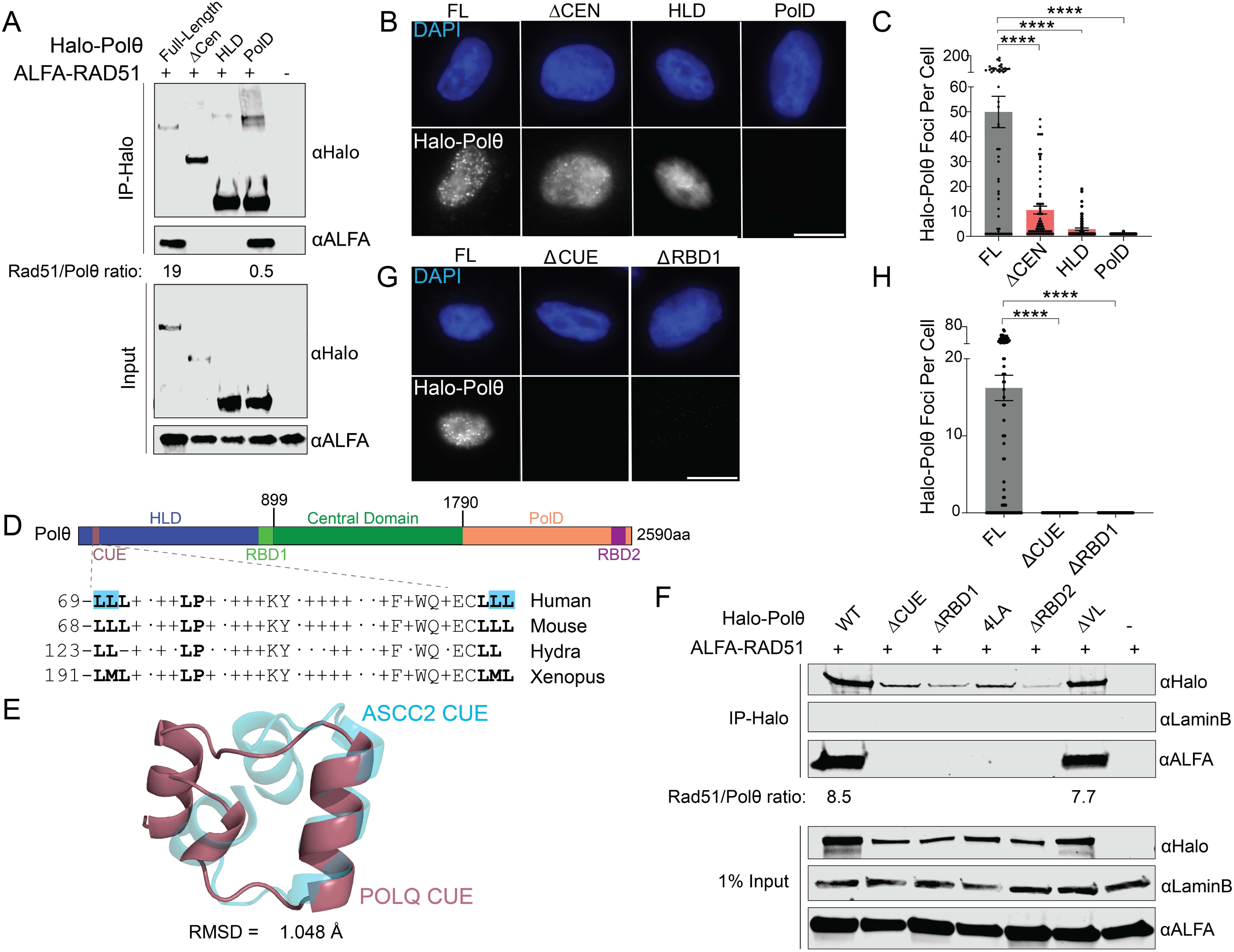
Structural features of Polθ required for RAD51 interaction and foci formation. (A) Nuclear extract co-IP of full-length (FL) Halo-tagged Polθ or Halo-Polθ domain mutants (∆CEN, HLD, or POLD) with ALFA-RAD51 in HEK293T cells. The bottom panels show 1% input. Quantification of ALFA-RAD51/Halo-Polθ ratio is indicated when interaction is detected. (B) Representative images of *TP53^−/-^* hTERT-RPE1 cells expressing Halo-Polθ alleles (FL, ΔCEN, HLD, POLD). (C) Quantification of nuclear Halo-Polθ foci. All constructs significantly differ from FL (****P < 0.0001, two-tailed Welch’s t-test). (D) Schematic of Polθ domain architecture: HLD, CUE, central domain, and PolD. RAD51-interacting subdomains: CUE, RBD1, and RBD2 are indicated. (E) Polθ CUE domain sequence alignment across species; asterisks indicate conserved residues. (F) Structural overlay of the predicted Polθ CUE domain (pink) with ASCC2 CUE domain (blue); RMSD = 1.048 Å. (G) Nuclear extract co-IP of Halo-Polθ mutant alleles with ALFA-RAD51 in HEK293T cells. Quantification of RAD51/Halo-Polθ ratio is indicated when interaction is detected. (H) Images of TP53^−/-^POLQ^−/-^ hTERT-RPE1 cells expressing Polθ mutants (FL, ΔCUE, ΔRBD1, ΔVL) induced by 16h of Dox and subsequent treatment with 100 ng/mL MMC for 6h. (G) Quantification of foci per cell. Significant reductions observed for all mutants (****P<0.0001, two-tailed Welch’s t-test). Data are mean ± SEM. Scale bars, 10 µm.

We also generated stable hTERT-RPE1 lines expressing the same Halo-Polθ transgenes under a Dox-inducible promoter and expression was confirmed by Western blot (Supplementary Figure 5A). Cells were induced with Dox for 16 hours, treated with MMC, and processed 6 hours later for immunofluorescence after cytosolic and nucleoplasmic pre-extraction. Halo-Polθ-FL formed robust MMC-induced chromatin foci, while the NLS-PolD constructs failed to form foci despite RAD51 interaction by co-IP (Figure 4B-C). The HLD and ΔCEN constructs showed significantly reduced and qualitatively distinct foci relative to FL. These findings indicate that helicase, central, and polymerase domains are all necessary for proper RAD51-colocalized Polθ foci formation after ICL induction.

Given that Polθ selectively interacts with ubiquitylated RAD51, we screened the *POLQ* coding sequence for putative ubiquitin binding domains. This analysis uncovered a conserved CUE (coupling of ubiquitin conjugation to endoplasmic reticulum degradation) domain within the N-terminal region of the Polθ-HLD (Figure 4D), showing significant structural similarity to the CUE domain in the known ubiquitin binding protein ASCC2 as defined by a low Root-Mean-Squared-Distance (RMSD) of ∼1 angstrom (Figure 4E)(*36*, *37*). Deletion of the Polθ-CUE domain (∆69-102) as well as alanine substitution of key di-leucine motifs (4LA, indicated by blue highlight in Figure 4D) abolished RAD51 interaction by co-IP analysis in HEK293T cells (Figure 4F). Furthermore, stable expression of Halo-Polθ^∆CUE^ in *TP53^−/-^POLQ^−/-^* hTERT-RPE1 cells demonstrated a loss of MMC-induced chromatin foci formation (Figure 4G-H).

A prior study identified a RAD51-binding region at the interface between the Polθ-HLD and CEN domains, which we refer to as RBD1 (aa846-895)(*1*). RBD1 overlaps the recently characterized Domain 5 (D5) within the Polθ-HLD, which undergoes substantial conformational rearrangement upon ssDNA binding—a prerequisite for microhomology annealing and TMEJ activity(*38*, *39*). We confirmed that Halo-Polθ^ΔRBD1^ lacks RAD51 interaction by nuclear co-IP in HEK293T cells and fails to form MMC-induced foci in *TP53^−/-^POLQ^−/-^* hTERT-RPE1 cells (Figures 4F-H). Structural modeling further suggests that the Polθ-HLD CUE and RBD1 domains are surface exposed and spatially proximal in the apo (non-ssDNA-bound) state (Supplementary Figure 5C), potentially facilitating cooperative engagement with ubiquitylated RAD51 nucleofilaments. Upon ssDNA binding, these domains reposition to opposite faces of the helicase (Supplementary Figure 5C), suggesting that RAD51 interaction occurs prior to loading of Polθ -HLD with ssDNA to perform microhomology (MH) annealing (*38*, *40*). Dimerization of the Polθ - HLD has been shown to be necessary for MH annealing and initiation of TMEJ (*38*, *39*). However, a Polθ mutant that disrupts the dimerization interface (Polθ ^∆VL^; ∆643-644) retains RAD51 binding (Figure 4F), consistent with the notion that Polθ interaction with RAD51 occurs in a monomeric, TMEJ inactive state.

To further delineate RAD51-binding determinants, particularly since we observed a PolD interaction with RAD51, we applied HADDOCK-based modeling using the Polθ-FL sequence(*41*, *42*). This analysis predicted a second high confidence RAD51-binding domain (RBD2; residues 2545-2575) within the Polθ-PolD (Supplemental Figure 5D). RBD2 deletion, similar to deletion of CUE and RBD1, abrogated RAD51 nuclear co-IP (Figure 4F).

Collectively, these data support a model in which Polθ engages ubiquitylated RAD51 filaments through multivalent interactions involving three distinct domains: the CUE domain within the HLD, RBD1 at the HLD-central domain interface, and RBD2 within the PolD. Polθ associates with RAD51 in a monomeric, apo state—prior to ssDNA loading and dimerization—suggesting that recruitment precedes TMEJ execution. Notably, Polθ engagement is increased when HR is stalled, as evidenced by a strand exchange-defective RAD51 mutant. These findings define a staged mechanism in which Polθ is recruited to stalled or unresolved RAD51 nucleofilaments via multivalent interactions with ubiquitylated RAD51 nucleofilaments, setting the stage for downstream RAD51 removal, ssDNA loading, and activation of TMEJ.

### MMC Induced Point Mutations are not Polθ Dependent

Having defined the molecular requirements for Polθ recruitment to ubiquitylated RAD51 filaments, we next sought to determine the functional consequences of TMEJ engagement during ICL repair. Specifically, we examined whether Polθ activity contributes to the mutational landscape arising from MMC-induced genotoxic stress. Towards this end, we performed whole genome sequencing (WGS, 45X) of five clones each from isogenic *TP53^−/-^* (i.e., POLQ “*WT*”) and *TP53^−/-^POLQ^−/-^*(i.e., “*POLQ^−/-^*”) hTERT-RPE1 cells treated with three sequential rounds of MMC (Figure 5A). Circos plots of single nucleotide variants (SNV), insertions/deletions (indel), and chromosomal rearrangements revealed widespread genomic perturbations – driven predominantly by SNVs – in MMC-treated clones, irrespective of *POLQ* status (Figure 5B).

**Figure 5.**
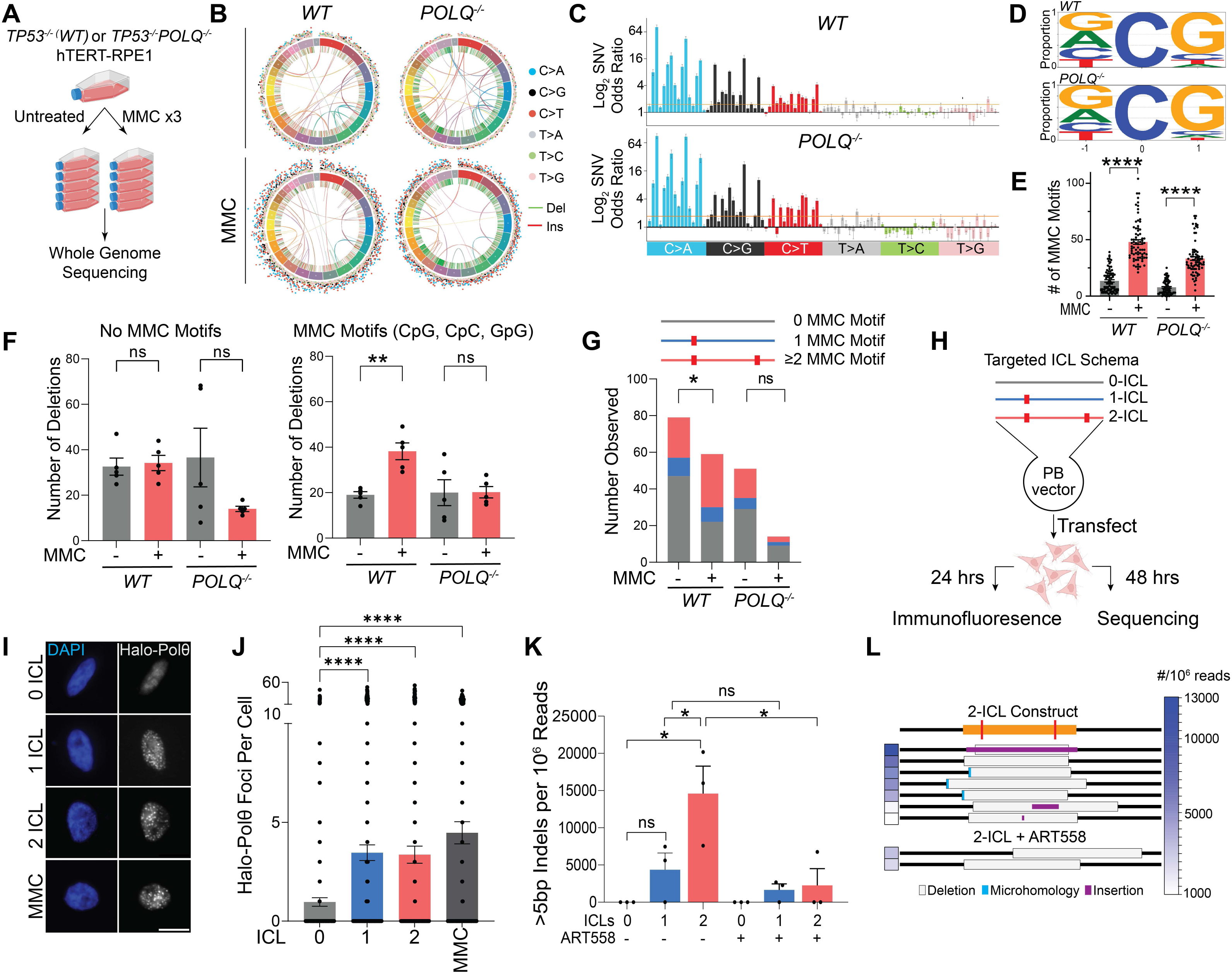
Polθ is essential for repair of proximal ICLs. (A) Schematic of mutagenesis workflow: sequential MMC treatment, clonal isolation, and WGS (∼45X) in *TP53^−/-^*(“WT”) and *TP53^−/-^POLQ^−/-^* (“*POLQ^−/-^*”) hTERT-RPE1 cell lines. (B) Circos plot representation of structural rearrangements, indels, and SNVs in untreated and MMC-treated *WT* and POLQ^−/-^ clones. Each Circos plot represents all 5 clones compared to the corresponding parental cell line. SNVs are shown as number per 5Mb interval and color coded based on mutation type. C>A/T/G SNVs are notably increased in both genotypes after MMC. (C) Odds ratio of MMC-treated to untreated COSMIC trinucleotide context single nucleotide variant (SNV) abundances in *WT* and *POLQ^−/-^* clones. (D) tSNE plot of COSMIC trinucleotide context SNV abundances demonstrates the distinct mutation signature observed after MMC treatment, and the similarity in mutation signatures between MMC-treated *WT* and *POLQ^−/-^* clones. (E) MMC-induced C>A/T/G mutations demonstrate an overrepresentation of CpG and CpC mutation contexts in both *WT* and *POLQ^−/-^* clones. (F) Quantification of deletions without (left) or with (right) MMC mutation-context motifs (CpG, CpC, GpG) (**P < 0.01, Mann-Whitney test). (G) Stacked histograms of microhomology-flanked deletions containing 0, 1, or >1 MMC motif, in untreated or MMC-treated *WT* and *POLQ^−/-^* clones (*P < 0.05, Fisher’s exact test). (H) Schematic of targeted ICL repair assay. Click chemistry-induced zero, single, or double interstrand crosslinked oligonucleotides were ligated into a Piggybac (PB) transposon vector, which was transfected into cells with Piggybac integrase. (I) Images of *TP53^−/-^POLQ^−/-^*hTERT-RPE1 cells expressing Dox-inducible Halo-Polθ 24 hours after transfection with 0-, 1-, or 2-ICL PB vectors. Also shown are cells treated with 100ng/mL MMC for 6 hours. (J) Quantification of Polθ foci (***P < 0.001, two-tailed Welch’s t-test). (K) TP53^−/-^ hTERT-RPE1 cells were harvested 48 hours after transfection with 0-, 1-, or 2-ICL PB vectors and PB integrase, either in the absence or presence of 10µM ART558 (Polθ inhibitor). NGS of amplicon PCR libraries across the crosslinked oligonucleotides were analyzed ScarMapperv2. Observed insertions/deletions (indels) >5bp per million reads are plotted for three biologically independent experiments (*P<0.05, two-tailed paired t-test). (L) Schematic of observed >5bp indels with abundance >1000 per 10^6^ reads in 2-ICL samples without or with ART558.

To further resolve the mutagenic processes involved, we applied trinucleotide context analysis, classifying each SNV by its base substitution type and flanking nucleotides, following previously defined frameworks for COSMIC mutational signatures (Supplementary Figure 6A)(*43*). Across both genotypes, the MMC-induced SNV profile resembled COSMIC signature SBS94, which is infrequently observed in human cancers and has unknown etiology (Supplementary Figure 6A) (*44*). Odds ratio analysis comparing treated versus untreated samples showed that the predominant substitutions in both genotypes were C>A, C>T, and C>G mutations (Figure 5C). Motif analysis revealed that in both genotypes these substitutions were significantly enriched at NpCpG and, to a lesser extent, NpCpC motifs (Figure 5D)—the known targets of MMC-induced interstrand and intrastrand crosslinks, respectively(*45*). Median SNV counts at these motifs were significantly induced by MMC in both *WT* and *POLQ^−/-^*clones (Figure 5E). Additionally, t-SNE projection, principal component, and bootstrap analyses showed no significant difference in trinucleotide spectra between genotypes following MMC treatment (Supplementary Figure 6B-D). These observations indicate that Polθ does not alter the dominant base substitution profile attributed to TLS during ICL repair in mammalian cells, similar to recent observations in *C. elegans*(*46*).

### Deletions Containing Multiple MMC Motifs are Polθ Dependent

Since SNV mutational spectra were indistinguishable between genotypes, we next focused on characterizing deletions based on the presence or absence of canonical MMC-reactive motifs (CpG, CpC, and GpG) within the deleted interval. Deletions lacking these motifs occurred at similar frequencies across all clones, regardless of genotype or treatment. In contrast, MMC-treated *WT* clones exhibited a significant increase in deletions containing MMC-reactive motifs, a pattern not observed in *POLQ^−/-^* clones (Figure 5F). To further define Polθ-specific events, we analyzed deletions flanked by >1bp microhomology (i.e., microhomology-associated deletions, “MHD”)—a genomic hallmark of TMEJ. MHDs containing 2 or more internal MMC-reactive motifs were significantly enriched in MMC-treated *WT* clones, but not in *POLQ^−/-^*clones (Figure 5G). These findings suggested that Polθ/TMEJ may be required to repair clustered ICL lesions, which may generate complex replication fork collapse events that are refractory to HR repair.

### Polθ is Required for Repair of Clustered ICLs

To directly test whether Polθ is required for resolving clustered ICLs, we designed double-stranded DNA oligonucleotides containing 0, 1, or 2 site-specific ICLs using a previously established click chemistry-based approach (Supplemental Figure 6E)(*47*). These duplex oligonucleotides were ligated into Piggybac transposon vectors and co-transfected with Piggybac integrase into *TP53^−/-^POLQ^−/-^*hTERT-RPE1 cells expressing doxycycline-inducible Halo-Polθ (Figure 5H). Robust nuclear Halo-Polθ foci were observed 24 hours post-transfection in response to the 1-ICL and 2-ICL constructs, but not the 0-ICL construct, mirroring the cellular response to MMC (Figure 5I-J).

To assess Polθ-dependent repair activity, the same ICL-containing constructs were transfected into *TP53^−/-^* hTERT-RPE1 cells in the presence or absence of the Polθ inhibitor ART558(*48*). After 48 hours, repair junctions were amplified and analyzed by high-throughput sequencing. Quantification of >5bp insertion/deletion (indel) events revealed a significant increase in indels in cells transfected with the 2-ICL construct, with a non-significant trend for the 1-ICL construct (Figure 5K). ART558 treatment abolished this increase, confirming the Polθ dependence of the indel genomic scars observed specifically in the repaired 2-ICL construct (Figure 5K).

Sequencing analysis of individual repair products from the 2-ICL construct revealed deletions spanning both ICLs, frequently accompanied by templated insertions and/or microhomology—hallmarks of TMEJ repair, consistent with resolution of DSBs arising from two-ended replication fork collapse at clustered ICLs (Figure 5L). In contrast, rare indels recovered in ART558-treated cells lacked these hallmarks (Figure 5L). These findings establish a specific requirement for Polθ in resolving clustered ICLs via TMEJ and support a model in which TMEJ functions as an essential salvage pathway when HR is unable to repair replication-associated multi-lesion damage.

## Discussion

Polθ-mediated end joining (TMEJ) has predominantly been viewed as a mutagenic backup pathway, engaged only when HR or non-homologous end joining (NHEJ) are absent or impaired(*1*, *2*, *9*, *10*). However, this view is difficult to reconcile with the deep evolutionary conservation of Polθ across most eukaryotic species, as these organisms typically retain functional canonical repair pathways. In this study, we demonstrate that TMEJ serves an essential and regulated role even in HR- and NHEJ-proficient mammalian cells, specifically by enabling the repair of clustered, replication-associated DNA lesions that cannot be resolved by HR alone. Rather than competing with HR, Polθ acts downstream of HR initiation and is recruited to stalled Rad51 nucleofilaments through multivalent interactions with ubiquitylated Rad51. In this poised state, Polθ is primed to engage when recombination intermediates fail to mature, positioning TMEJ as a hierarchical fail-safe that preserves genome integrity under replication stress.

In the context of ICL repair, we define a hierarchical repair cascade in S phase that culminates in Polθ recruitment to sites of replication fork collapse. This process depends on sequential upstream events, including p97-dependent CMG helicase unloading, FANCD2- and SLX4-mediated ICL unhooking, and BRCA2-dependent RAD51 filament formation (Figure 6). Crucially, Polθ is recruited only to a subset of mature RAD51 nucleofilaments that have undergone ubiquitylation by FBXO5 or FBH1. This selective targeting requires multivalent interactions with ubiquitylated RAD51, mediated in part by a previously uncharacterized CUE ubiquitin-binding domain within the Polθ helicase-like domain. Disruption of this domain, or inhibition of RAD51 ubiquitylation, impairs Polθ foci formation and its association with RAD51 filaments. This specificity ensures that Polθ recruitment does not interfere with earlier HR steps such as end resection and RAD51 loading, and instead is recruited to mature HR intermediates. It also offers a mechanistic explanation for the increased reliance of HR-deficient cells on PLK1- and RHNO1-dependent Polθ recruitment in M phase, where the S-phase RAD51-dependent recruitment pathway is impaired(*12*, *18*, *49*, *50*).

**Figure 6.**
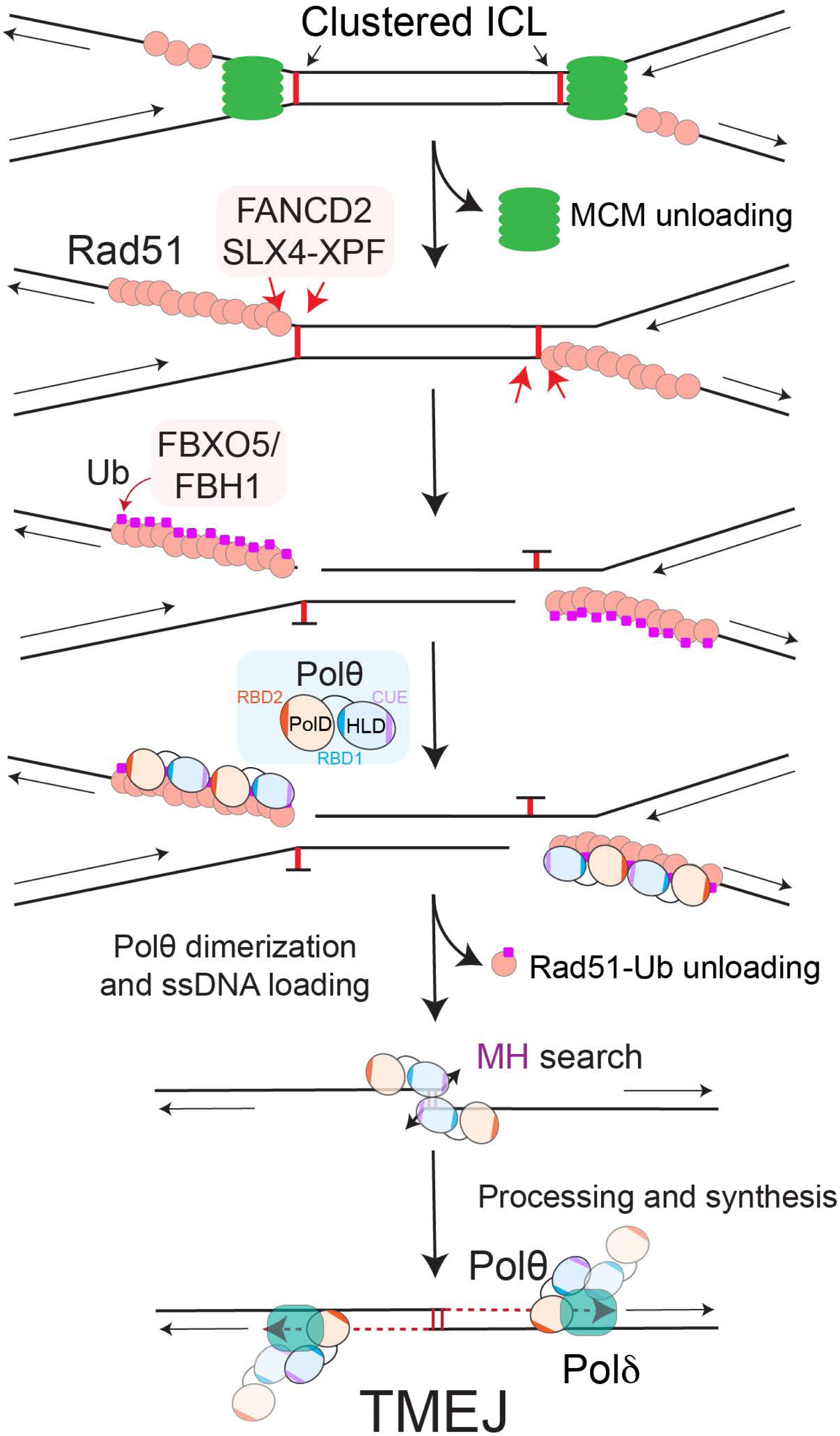
Model of regulated Polθ recruitment to resolve clustered ICLs via TMEJ. Following MCM unloading and nucleolytic unhooking of clustered interstrand crosslinks, bidirectional replication fork collapse can generate a two-ended DSB that has been loaded with RAD51 but is incompatible with HR due to lack of a sister chromatid spanning the break site. At these persistent RAD51 nucleofilaments, ubiquitylation by FBXO5 and FBH1 licenses Polε recruitment via multivalent interactions, including a CUE domain and two additional RAD51 binding motifs. The mechanism of RAD51 unloading in this context remains unresolved and may require additional factors. Additionally, after RAD51 is unloaded Polε must dimerize and be loaded with ssDNA to initiate TMEJ, a process that might require the ATPase activity of the Polε-HLD. This regulated, hierarchical deployment of Polε likely extends to other lesions that induce convergent fork collapse in the absence of an intervening origin, defining a broader role for TMEJ in resolving HR-incompatible replication-associated DNA damage.

Although Polθ can function as a TLS polymerase *in vitro*, our genomic scar analyses found no evidence of such a role in cells *in vivo* during ICL-induced mutagenesis, consistent with another recent study that evaluated Polθ-dependent mutation scars using a distinct ICL substrate in *C. elegans*(*46*, *51*). In contrast, we uncovered a specific requirement for Polθ/TMEJ in the repair of clustered ICLs using both genome-wide scar analyses and an engineered site-specific ICL substrate. While Polθ/TMEJ-dependent microhomology-associated deletions containing multiple MMC-reactive motifs accounted for only ∼1% of total ICL-induced mutations, these events likely underlie the heightened sensitivity of Polθ-deficient cells to MMC treatment.

The precise topology of replication fork collapse at clustered ICLs, and why these lesions might be refractory to HR, remains incompletely understood. Recent studies indicate that single-strand nicks encountered during replication can produce either one-ended and two-ended DSBs, depending on whether the lesion is on the leading or lagging strand, respectively(*3*, *52*, *53*). In both scenarios, the availability of an intact sister chromatid generally permits efficient HR-mediated repair. In contrast, clustered ICLs may stall converging replication forks on both the leading and lagging strands, obstructing fork progression through the intervening region (Figure 6). After nucleolytic processing, this configuration could yield two-ended DSBs within the parental duplex DNA, without a continuous sister chromatid spanning the break, thereby precluding effective strand invasion and completion of HR. The resulting DNA structures—RAD51-loaded ssDNA filaments at both ends—may not be amenable to resolution by error-free recombination pathways (Figure 6). Recruitment of Polθ to these sites may thus represent a salvage pathway, leveraging the RAD51-coated ssDNA as a substrate for TMEJ engagement. This mechanism may preserve genetic information by limiting large deletions or chromosomal rearrangements, albeit at the cost of TMEJ associated small indels.

Such co-incident, two-ended replication fork collapse events may be more common under conditions of heightened replication stress and are likely exacerbated in cancer cells with oncogene-induced replication stress. The essential requirement for Polθ/TMEJ in these contexts may explain the broad synthetic lethal landscape associated with Polθ dependency, its frequent overexpression in diverse cancers, and its synergy with ATR inhibition(*5*, *10*, *20*).

Polθ’s selective binding to ubiquitylated RAD51 nucleofilaments through multivalent interactions— including the CUE, RBD1, and RBD2 domains—provides a molecular mechanism to discriminate DNA-bound RAD51 from soluble RAD51 pools. Monomeric Polθ engages RAD51 via these domains but is structurally incompatible with TMEJ, which requires Polθ dimerization to coordinate ssDNA binding and microhomology search(*39*). RBD1 overlaps with the D5 domain of the Polθ-HLD, which undergoes a major conformational change upon ssDNA binding—a transition likely incompatible with stable RAD51 binding(*38*, *39*). We propose that Polθ recognizes stalled or persistent RAD51 filaments but requires RAD51 unloading—potentially mediated by FIGNL1 or other anti-recombinases—for TMEJ to proceed. The transition from HR to TMEJ may thus involve coordinated regulation by Polθ and additional HR factors—a poorly understood process that warrants further investigation.

Understanding this regulated transition from HR to TMEJ helps resolve a longstanding paradox: how a mutagenic repair pathway like TMEJ can nonetheless promote genome stability and be evolutionarily conserved. Polθ’s highly restricted deployment ensures it is activated only when HR fails, enabling repair of otherwise irrecoverable lesions while minimizing mutagenic risk. Our findings align with prior studies in *C. elegans*, where Polθ deficiency leads to large deletions at G4 quadruplex-associated replication fork collapse—possibly reflecting an inability to salvage RAD51-loaded nucleofilaments when HR is unsuccessful(*21*, *54*).

These insights also have important therapeutic implications for the potential application of Polθ inhibitors in HR proficient cancers. Our findings suggest that Polθ inhibition may be effective in tumors with RAD51 overexpression, elevated replication stress, and during therapy with replication stress inducers. Supporting this notion, Polθ has been shown to promote survival following high-LET radiotherapy, which is known to generate clustered DNA damage(*55*). These contexts may define a therapeutic niche for Polθ inhibitors, whose rational deployment, particularly in HR proficient cancers, will benefit from further insight into the molecular logic governing Polθ ‘s recruitment and activity.

## Acknowledgments

We are grateful to members of the Gupta lab, Richard Wood, and Neil Johnson for helpful discussions, and to Pablo Ariel for microscopy expertise.

## Funding

This work was supported by grants from the NIH/NCI P01 CA247773 (to G.P.G, E.R., D.A.R, S.D., and R.D.W.), R37 CA227837 (to G.P.G.), BCRF (to G.P.G.), P50 CA058223 (to G.P.G.), Yang Biomedical Scholar Award (to G.P.G.), UNC TNBC Center (to G.P.G.), 1T32GM12274 (to C.M.S.), and R50CA233185 (to B.E.E.). G.P.G. is a recipient of the Burroughs Wellcome Career Award for Medical Scientists. Core Facility Services (Microscopy Services Laboratory and Flow Cytometry Core) are supported by the Cancer Center Support Grant (P30 CA016086) to the UNC Lineberger Comprehensive Cancer Center.

## Competing interests

G.P.G. receives patent licensing fees from and has equity in Naveris, Inc, and is the recipient of research funding from Breakpoint Therapeutics and Merck.

