## Supplementary material for "Hierarchical Coordination of Polymerase Theta and RAD51 Resolves Clustered Replication Fork Collapse": Methods and Supplemental Figures

### Materials and Methods

#### Cell Culture

Cells were grown in a humidified incubator at 37 °C and 5% $CO_2$. HEK293T (ATCC) cells and hTERT RPE1 (ATCC) cells were grown in DMEM, high glucose (Corning, 10-013-CV ) supplemented with 10% fetal calf serum (Corning, 35-015-CV) or Hyclone bovine growth serum (Cytiva, SH30541.03). Cells were split at approximately 80% confluence using 0.05% Trypsin EDTA Solution (Gibco). For cells containing puromycin-resistant constructs, 1 ug/mL Puromycin (ThermoScientific) was included in the growth media. All cell lines were routinely screened for mycoplasma contamination using MycoStrip™Mycoplasma Contamination Kit (Invivogen, rep-mysnc-100) according to the manufacturer's protocol.

#### Polymerase Theta Expression Constructs

All expression constructs were verified by sequencing (Plasmidsaurus) before use in cell line construction. A retroviral expression vector containing Halo-tagged polymerase theta (Halo-Polθ) cDNA was obtained from Genewiz (30-330513735). The Halo-Polθ cDNA from pBABE-Puro+Halo-Polθ was cloned into PB-TA-ERP2 to create a full-length, doxycycline inducible Halo-Polθ expression vector (pGGL-65). Cloning was done by In-Fusion (TaKaRa) using PrimeStar GXL Premix polymerase (TaKaRa, R051A L3000001). All constructs were transformed into Stbl3™ cells (Invitrogen, C737303).

#### Cell Line Derivation

Cell lines were electroporated using the Invitrogen Neon Transfection System with a setting of 1350 V, 20 milli-second pulse, 2x. 200,000 to 300,000 cells were used with the 10 µL tip for sgRNA. hTERT RPE1 *TP53*$^{-/-}$ and hTERT RPE1 *TP53*$^{-/-}$ *POLQ*$^{-/-}$ cells were previously described (*1*).

Retrovirus packaged from the pBABE-Puro+Halo-Polθ construct was used to transduce hTERT RPE1 cells to derive the hTERT RPE1 Halo-Polθ C26 cell line. Following transduction, cells were selected with puromycin and sorted for Halo expression. Halo-Polθ expression was validated by Western blot.

To construct cell lines based on PB-TA-ERP2 constructs including Halo-Polθ$^{FL}$, Halo-Polθ$^{HLD}$, Halo-Polθ $^{\Delta CEN}$, and Halo-Polθ $^{\Delta VL}$ , 800,000 cells were suspended in 100 µL of Buffer R containing 200 ng of a Piggybac integrase expression vector (System Biosciences) plus 2 µg of a PB-TA-ERP2 construct and electroporated using the Neon Transfection System (Invitrogen). Cells were immediately seeded into a 15 cm dish in 25 mL of media without selection. After 48 hours incubation, 1 µg/mL puromycin was added to the media.

To construct cel lines based on PB-TA-ERP2 constructs including Halo-Polθ$^{NLS\text{-}PolD}$, Halo-Polθ$^{\Delta RBD1}$, and Halo- Polθ$^{DCUE}$ , 250,000 cells were seeded in a 6 well dish 6 hours prior to transfection. Cells were transfected following the Lipofectamine 3000 (Thermo Fisher) reagent protocol, using 1 µg of PB-TA-ERP2 construct and 0.1 µg of a Piggybac integrase expression vector (System Biosciences). After 48 hours incubation, 1 µg/mL puromycin was added to the media.

Activity of the exogenous Polθ was measured using the PathSig assay as previously described (*2*)

#### Colony formation assay

hTERT RPE1 *TP53*$^{-/-}$ and hTERT RPE1 *TP53*$^{-/-}$ *POLQ*$^{-/-}$ cells were seeded in triplicate for each condition, at 500 cells (0, 5, and 10 ng/mL Mitomycin C (MMC, Millipore-Sigma, M4287)) or 2000 cells (15, and 20ng/mL MMC) per 10 cm dish in 10 mL media. The cells were allowed to grow for 14 days without changing the media. At this time the media was removed, and colonies were stained with a solution of 0.05% crystal violet (Sigma C0775) in 40% methanol and manually counted.

**Metaphase spreads**

hTERT RPE1 *TP53*$^{-/-}$ or hTERT RPE1 *TP53*$^{-/-}$ *POLQ*$^{-/-}$ cells were seeded 48-hours prior to harvesting at a density to allow logarithmic growth at time of harvest. 10 ng/mL MMC was added to the media 15 hours prior to harvesting. Metaphase spreads were prepared as previously described(*3*). Metaphases were imaged using a Keyence BZ-X810 microscope with a 100x objective.

**Lentiviral Production and Cell Transduction**

Five million cells were infected at a 0.5 multiplicity of infection for 24 hours in media containing 4 µg/mL hexadimethrine bromide (Millipore-Sigma). Following infection, the cells were passaged and seeded at a density to obtain ≤ 80% confluent after 48 hours. After 48 hours, the cells were trypsinized and suspended into a single cell suspension, then sorted for mVenus expression. Five million cells were recovered from the sort and were seeded in T175 flasks at ≤ 1 million cells and allowed to recover for 24 hours. For all subsequent cell passages, five million cells were maintained in culture and seeded at one million cells per T175 flask.

**Cas9 Activation and Drug Treatment**

Twenty-four hours after cell sorting, 5 µg/mL Shield-1 (Aobious, AOB1848) was added to stabilize Cas9. The cells were left for 24 hours in Shield-1 then passaged and seeded at 1-million cells per T175 flask. Extra cells were washed with Dulbecco's phosphate buffered saline (PBS) and stored as a frozen pellet without buffer at -80 °C. These were used as the sample control for essential genes. Six to eight hours after seeding, 20 ng/mL MMC was added to the media. The MMC was removed after 15 hours. Forty-eight hours after seeding the cells were again passaged. This was done for a total of three MMC treatments. Forty-eight hours after the last treatment the cells were placed into single cell suspension, washed with PBS, and frozen at -80 °C.

**Whole Genome Sequencing Drug Treatment and Library Preparation**

One million hTERT RPE1 *TP53*$^{-/-}$ or hTERT RPE1 *TP53*$^{-/-}$ *POLQ*$^{-/-}$ cells were each seeded into a T175 flask. The next day 20 ng/mL MMC was added to each flask. After 15 hours the media was changed. The cells were allowed to recover for 48 hours at which time they were passaged, and 1 million cells were reseeded into a T175 flask. This was repeated and the cells were treated with MMC and allowed to recover a total of 3 times. Following the final recovery, 5 clones were selected from the treated and untreated cell populations. These were expanded for genomic DNA purification using the Qiagen DNeasy kit. Next Generation Sequencing libraries were prepared from the cell populations, five clones from each of the untreated populations, and five clones from each of the treated populations by tagmentation using the Illumina Nextera FLEX kit (20018704). Library preparation was according to the manufacturer's protocol. Libraries were quantified, pooled, and sequenced at 2x150 cycles on three lanes of a Novoseq6000 S4 (Novogene).

**Whole Genome Sequence Analysis**

Adapter sequences were trimmed from each set of the FASTQ files using Atropos. After demultiplexing, the individual FASTQ files for each lane were aligned to the GCRh38 human reference genome using BWA with default settings. Read groups were added to the BAM files using picard tools AddOrReplaceReadGroups. After adding read groups, the BAM files were deduplicated using Samtools MarkDup set to remove the PCR duplicates. Finally, the BAM files from each Novoseq6000 lane were merged using Samtools merge. Variant calling in post-MMC treated daughter subclones were called as previously described (*4*) using CaVEMan (v.1.13.15) (*5*), Pindel (v.3.2.0) (*6*) and BRASS (v6.3.4). Specifically, for single-nucleotide variant (SNV) calls by CaVEMan, we used CLPM = = 0 and ASMD ≥ 140. To reduce false positive calls by Pindel we used QUAL ≥ 250 and REP < 10. Cell clones with an average variant allele frequency of <0.4 were designated as polyclonal and excluded from all subsequent quantitative analyses. De novo substitutions and InDels in subclones were obtained by subtracting variants in respective parental clones, and by removing mutations shared among subclones.

**Knockdowns by siRNA**
Cells were plated 24 hours before treatment at 200,000 cells per well in a 6-well dish. siRNAs targeting FANCC, FANCD2, SLX4, ERCC4, RFWD3, FBXO5, RAD51, BRCA2, or no target control (Horizon Discovery), were added per protocol using Lipofectamine RNAiMAX Transfection Reagent (Invitrogen, 13778075). 24 hours after the first siRNA treatment, this was repeated. 48-hours following initial siRNA treatment, cells were replated for RT-PCR or immunofluorescence. Knockdown efficiency was determined through RT-PCR.

**RAD51 mutant generation and transfection**
Gene blocks (gBlocks) were designed and ordered through Integrated DNA Technologies (Integrated DNA Technologies). gBlocks generated included RAD51 wildtype, RAD51 K57/58/64/107/156R (5KR), RAD51 R130/303A and K313A (ii3A), and a RAD515KR/ii3A double mutant. Each sequence was also designed to be resistant to siRNA targeting endogenous RAD51, through silent mutations in the siRNA target sequence, and an ALFA tag on the N-terminal of RAD51. Using Gateway cloning (Invitrogen, 11791043 and 11789100), gBlocks were cloned into a pDONR223 backbone through BP reaction, and a PB-TA-ERP2 destination vector (PB-TA-ERP2 was a gift from Knut Woltjen (Addgene plasmid # 80477; http://n2t.net/addgene:80477; RRID:Addgene_80477)) through LR reaction. DH5a E. coli was used for transformation and DNA extraction was done using Qiagen QIAprep Spin Miniprep Kit. Cells were transfeted using Lipofectamine 3000 Transfection Reagent (Invitrogen, L3000001) followed by selection with media containing 1 ug/mL puromycin. Expression of ALFA-tagged RAD51 constructs was confirmed by Western blot. Following knockdown of siRAD51 in ALFA-RAD51 containing cell lines, 1 ng/mL (immunofluorescence) or 100 ng/mL (Western blot) Doxycycline (Sigma-Aldrich) was added to media to induce expression of RAD51 siRNA resistant mutants. At 16 hours, respective drug treatment followed by immunofluorescence was performed. Expression was confirmed using western blot.

**Immunofluorescence**
Cells were plated on glass coverslips (VWR VistaVision No. 1.5) in 6-well dishes at 200,000 cells/well in 2 mL of media. 24-hours after plating, respective drug treatments and HaloTag Ligand JF669 (Janelia Farm) or JFX646 (Promega, GA1121) were added, and cells were incubated for 6 hours (37ºC, 5% $CO_2$). When applicable, EdU was added 30 minutes before fixation of slides to label S phase cells (BaseClick, Millipore-Sigma, BCK-EDU488). Cells were permeabilized with CSK buffer (10 mM HEPES, 300 mM Sucrose, 100 mM NaCl, 3 mM $MgCl_2$, and 0.5% Triton X-100, pH 7.4) for 4 minutes at room temperature and fixed with 4% paraformaldehyde diluted in PBS for 10 minutes. Edu labeling kit was used when applicable (BaseClick). Slides were blocked for 1 hour with 5% goat serum (Jackson ImmunoResearch Labs, 005-000-001) in PBS at room temperature. Primary antibodies were diluted in 5% goat serum, (mouse anti-RAD51 14B5 at 1:500, rabbit anti-RAD51 D410B 1:500, rabbit anti-ALFA at 1:2,000, mouse anti-PCNA P10 at 1:500) and slides were incubated for 30 minutes at room temperature. Slides were washed two times with PBS. Secondary antibodies diluted in 5% goat serum at 1:500 and incubated on slides for 30 minutes. Slides were washed two times with PBS. DAPI (Thermo Fisher Scientific, 62248) was diluted at 1:10,000 in PBS and incubated on slides for one minute. Slides were washed with PBS and mounted with ProLong Gold Mounting Media on microscope slides (VistaVision$^{TM}$, VWR) and allowed to dry overnight. Slides were sealed with nail polish and imaged approximately 24 hours after mounting. Slides were protected from light throughout the experiment.

**Single-Molecule Localization Microscopy (STORM)**

Sample preparation, imaging and data analysis for STORM were performed in triplicate as described previously (*7, 8*). Briefly, hTERT RPE1 cells expressing Halo-Polθ were trypsinized and seeded on glass coverslips (Fisher Scientific, #12-548-B) in six-well plates in low density. Treatment with 100 nM Camptothecin (CPT) (Santa Cruz/ChemCruz, sc-200871B) (or DMSO) was performed directly on cells on coverslips for 1 hour. To detect cells in S phase, cells were pulse-labeled with 10 mM EdU (ThermoFisher #A10044) 30 minutes into the treatment. Cells were then allowed to recover for either 0 or 4 hours, after which they were permeabilized with 0.5% Triton X-100 in ice-cold CSK buffer at room temperature for 3 minutes and fixed with 4% paraformaldehyde (Electron Microscopy Sciences #15714) in RT for 15 minutes. Following fixation, cells were washed twice with PBS and blocked with blocking buffer (2% glycine, 2% BSA, 0.2% gelatin, and 50 mM $NH_4Cl$ in PBS). After fixation, EdU was tagged with CF® Dye Picolyl Azide CF488A (Biotium #92187) through click chemistry (Click-iT chemistry, ThermoFisher, #C10640), followed by a 30-minute blocking step. Cells were then stained for RAD51 (Abcam, #GR3270300-14, 1:2000 dilution) for 1 hour in blocking buffer and after that with a secondary Alexa Fluor 647-conjugated antibody (Invitrogen, #A-21245, 1:10000 dilution) for 30 minutes. Immediately prior to imaging, coverslips were mounted on glass slides to create a chamber, which was filled with imaging buffer (PBS with 100 mM mercaptoethylamine (Fisher Scientific, #BP2664100), 1 mg $ml^{-1}$ glucose oxidase (Sigma, #G2133), 0.02 mg $ml^{-1}$ catalase (Sigma, #C3155) and 10% glucose (Sigma, #G8270)).

STORM imaging was performed on a custom-built optical imaging platform described in detail elsewhere (*7, 8*). 2000 frames were acquired per field of view at a frequency of 33 Hz, with each replicate comprising at least 6 different fields of view. Localization of single molecules was performed by 2D-Gaussian multi-PSF fitting, as described previously (*7*). Correlation profiles were generated as a function of the pair-wise distances between targets and then fitted to a Gaussian model of a target species with itself (auto-pair correlation) or another target (cross-pair correlation). The normalized amplitude of the cross-pair correlation indicates the extent of enrichment between both targets, in this case RAD51 and Halo-Polθ.

**RT-qPCR**

RNA samples were extracted using NucleoSpin RNA kit (Takara bio, 740955.250). The RNA concentrations of each sample were determined by Qubit RNA Broad Range Assay Kit (ThermoFisher Scientific, Q10211). 1 µg RNA was added to each cDNA synthesis reaction cDNA synthesis using Maxima First Strand cDNA Synthesis Kit for RT-qPCR (ThermoFisher Scientific, K1671) and cDNA concentration was measured using Qubit™ dsDNA BR Assay Kit (ThermoFisher Scientific, Q32853). 1 to 2 ng of cDNA from each sample was used in qPCR reactions using IG SYBR Green qPCR 2X Master Mix (Intact Genomics, 3357) in a 384-well plate format. QuantStudio 6 Flex Real-Time PCR System (Applied Biosystems) was used to perform qPCR and the data was analyzed by QuantStudio Real-Time PCR Software. The primer sets used in quantifying each gene can be found in the Supplemental Table.

**Cell Fractionation**

Fractionation was done by the method of Mendez et. al., 2000 (*9*), with the following modifications: Buffers A and B contained 1x Halt Protease Inhibitor cocktail (ThermoFisher, # 78438), 0.1% sodium azide, 25 mM sodium fluoride, and 5 mM β-glycerophosphate disodium salt pentahydrate. The buffers were formulated, filter sterilized, and stored at 4 ºC until use. Two to three million cells per condition were seeded into 15 cm dishes with the appropriate amount of doxycycline to induce expression. Twenty-four hours later the cells were trypsinized and lysed in Buffer A at a concentration of 15,000 cells per µL. The concentration was adjusted at each step to maintain the starting concentration. For Western blots, the insoluble nuclear fraction was suspended in 25% final volume of TE(8) + 4 mM

MgCl2 + 25-50 units of Benzonase® Nuclease (MilliPore-Sigma 70664-3). This was allowed to digest for 15 minutes at room temperature then enough 7 M Urea Buffer + 1 % SDS was added to bring the concentration to the desired amount (7M Urea, 2M Thiourea, 30 mM Tris-Cl pH 7.5).

**Western Blot**

Unless otherwise noted, 125,000 – 150,000 cells per lane were loaded per lane of a 26-well Bio-Rad Criterion 4 –15% precast gel (Bio-Rad 5671085). Electrophoresis was done at 200 volts until the dye front ran off the bottom. Proteins were transferred onto nitrocellulose (LICORbio #962-31092) membranes using wet transfer at 60 volts for 75 minutes. The membrane was blocked in tris-buffered saline pH 7.5 (TBS) containing 1% Cold Water Fish Skin Gelatin (Millipore-Sigma) for 1 hour rocking at room temperature and then incubated overnight at 4°C with the indicated primary antibody diluted in TBS+0.1% Tween-20 (TBST) containing 5% bovine serum albumin and 0.05% sodium azide. Subsequently, the membranes were washed 3 x 10 minutes in TBST. Membranes were incubated in secondary antibodies diluted 1:10,000 in 5% Bio-Rad Blotting-Grade Blocker dissolved in TBST for 2 hours at room temperature. Membranes were then washed again 3 x 10 min in TBST before a final wash in TBS for 3 minutes. Membranes were imaged using LICORbio Odyssey imager. Images were analyzed with Empiria Studio® Software (LICORbio).

**Co-Immunoprecipitation**

Cells were seeded into 10 cm dishes and transfected with 10 μg of Halo-Polθ expression plasmid and, if co-transfected, 3 μg of ALFA-RAD51 expression plasmid. Plasmid expression was induced by adding 100 ng/mL doxycycline. Cells were harvested 16-24 hours later for immunoprecipitation (IP).
During cell fractionation, cytoplasm was removed as described above. The nuclear pellet was then resuspended in 200 μL Pierce RIPA buffer (ThermoFisher, #89900) containing 7,000 units of DNase I, 2.5 mM $MgCl_2$, and 1× Halt Protease Inhibitor Cocktail, followed by 10 min incubation on ice. Samples were then sonicated at 50% amplitude for three cycles of 15 sec on and 10 sec off and cleared by centrifugation (3,000 × g, 4°C). Immunoprecipitation was done with ChromoTek Halo-Trap Magnetic Agarose beads (Proteintech, # OTMA) according to manufactures' recommendation with the following changes. 15 μL of beads were used for binding step, bead elution was performed with 50 μL Laemmli sample buffer (120 mM Tris-HCl, pH 6.8, 20% glycerol, 4% SDS, 10% β-Mercaptoethanol), and samples were boiled at 95°C for 5 minutes prior to loading on a Criterion TGX precast gel (Bio-Rad, #5671084). Input and bound fractions were analyzed by SDS-PAGE and Western blot as described above.

**Crosslinked Oligonucleotides; Synthesis and Purification**

Standard DNA phosphoramidites, solid supports (controlled pore glass) and additional reagents were purchased from Link Technologies, Glen research, Jena Bioscience, Berry Associates and Applied Biosystems. All oligonucleotides were synthesized on an Applied Biosystems 394 automated DNA/ RNA synthesizer using a standard 1.0 μm phosphoramidite cycle of acid-catalyzed detritylation, coupling, capping, and iodine oxidation. TCA (3% in dichloromethane), 1*H*-tetrazole (0.45 M in acetonitrile), Cap A (10% acetic anhydride, 10% lutidine and 80% tetrahydrofuran) / Cap B (16% N-methylimidazole in tetrahydrofuran) and iodine (0.02 M in tetrahydrofuran, pyridine and water) were used. Pre-packed nucleoside SynBaseTM CPG 1000/110 (Link Technologies) resins were used and β-cyanoethyl protected phosphoramidites (dA-bz, dG-ib, dC-bz and dT where bz = benzoyl and ib = *iso*-butyryl, Sigma-Aldrich) were dissolved in anhydrous acetonitrile (0.1 M) immediately prior to use. The coupling time for normal A, G, C, and T monomers was 60 s and the coupling time for 5'-(4,4'-Dimethoxytrityl)-N6-benzoyl-N8-[6-(trifluoroacetylamino)-hex-1-yl]-8-amino-2'-deoxythymidine, 3'-[(2-cyanoethyl)- (N,N-diisopropyl)]-phosphoramidite (amino modified C6 dT) (Link) and 5'-Dimethoxytrityl-5-(octa-1,7-diynyl)-2'-deoxyuridine, 3'-[(2-cyanoethyl)-(N,N-diisopropyl)]-phosphoramidite (alkyne) (Glen research) was extended to 840 s. Stepwise coupling efficiencies were determined by automated trityl cation conductivity monitoring and were >98% in all cases.

**Selective β-cyanoethyl Removal**
DNA strand bearing the amines were treated on-column with 20% diethylamine in anhydrous acetonitrile for 20 min at room temperature. The resin was then washed with acetonitrile (3 x 1 mL) and dried with argon.

**DNA Deprotection**
DNA was cleaved from solid support and deprotected by exposure to a concentrated solution of aqueous ammonia in a sealed vial for 5 h at 55 °C. After drying *in vacuo*, oligonucleotides were dissolved in water and subject to further purification.

**RP-HPLC Purification**
Oligonucleotides were purified using a Gilson reverse-phase high performance liquid chromatography (RP HPLC) system with an ACE® C8 column (particle size: 10 µm, pore size: 100 Å, column dimensions: 10 mm x 250 mm) with a gradient of buffer A (0.1 M triethylammonium bicarbonate (TEAB), pH 7.5) to buffer B (0.1 M TEAB, pH 7.5 containing 50% v/v MeCN) and flow rate of 4 mL/min. The gradient was increased from 0% to 100% buffer B over 30 minutes (Condition A). Elution was monitored by UV absorbance at 295 nm. After HPLC purification, oligonucleotides were freeze dried then dissolved in water without the need for desalting.

**Oligonucleotide mass spectrometry**
All DNA were characterized by negative-mode electrospray using a UPLC-MS Waters XEVO G2-QTOF mass spectrometer and an Acquity UPLC system with a BEH C18 1.7 µm column (Waters). A gradient of methanol in triethylamine (TEA) and hexafluoroisopropanol (HFIP) was used (buffer A, 8.6 mM TEA, 200 mM HFIP in 5% methanol/water (v/v); buffer B, 20% v/v buffer A in methanol). Buffer B was increased from 0–70% over 7.5 min or 15–30% over 12.5 min for normal oligonucleotides and 50–100% over 7.5 min for hydrophobic oligonucleotides. The flow rate was set to 0.2 mL/min. Raw data were processed and deconvoluted using the deconvolution software MassLynx v4.1.

**Post-synthetic oligonucleotide modification with azide active ester**
To synthesize the azide oligonucleotide, 6-Azidohexanoic acid NHS ester (1 mg) was added in DMSO (40 µL) post-synthetically to the freeze-dried amino-modified oligonucleotide (oligonucleotide modified with amino C6 dT) (200 nm) in 0.5 M $Na_2CO_3$/$NaHCO_3$ buffer, pH 8.75 (60 µL). After 4 h at room temperature the fully-labelled oligonucleotide was purified by reversed-phase HPLC.

**Click reaction to synthesise the crosslinked ICLs substrates***(10-12)*
The alkyne and azide single strands oligonucleotides (40 nm each) were dissolved in 0.2 M NaCl (400 µL), annealed by heating at 95°C for 5 min then gradually cooled to room temperature over 2 h. The solution was flushed with a stream of argon and to this was added an aqueous solution (0.2 M NaCl) of the $Cu^1$ click catalyst solution. $Cu^1$ click catalyst solution was prepared from tris(3-hydroxypropyltriazolylmethyl) amine ligand (*13*) (5 mg), $CuSO_4$ (20 µL, 100 mM), in an aqueous solution of sodium ascorbate (40 µL, 500 mM) then was thoroughly degassed using argon. The click mixture was thoroughly degassed using argon then left at room temperature for 2 h. Reactions were desalted using NAP-25 (GE Healthcare). Formamide (460 µL) was added and the samples analysed and purified using denaturing 8 % PAGE Urea gels (Figure S6). The gel was visualised by placing it on top of a TLC plate and G:Box then using a UV lamp at 254 nm to cut bands. Bands corresponding to the assembled ICLs were excised and extracted from the gel using the ‘crush and soak method’. The excised polyacrylamide pieces were broken down into small pieces then suspended in distilled water (25 mL). The suspension was shaken at 37 °C for 18 h then filtered through a plug of cotton wool. The filtrate was concentrated to approximately 2 mL then desalted through NAP-25 then NAP-10 columns. The desalted eluent was lyophilized prior to use. The pure cross-linked ICLs were characterised and confirmed by mass spectrometry and lyophilised prior to use.

**Targeted Interstrand Cross Link Repair**

Duplexed oligonucleotides were ligated into pGGL-31 that had been double digested with XhoI (NEB, R0146S) + SpeI-HF (NSB, R3133S) using a 5:1 molar ratio of oligonucleotide to vector. 170 fM of linear plasmid was ligated in a 100 µL reaction overnight at 18 ºC. Each 100 µL ligation was then mixed with 20 µL of 10x NEB Buffer 4 + 20 µL of ATP + 55 µL dd$H_2O$ + 5 µL of Exonuclease V (RecBCD) (New England Biolabs) and incubated for 2 hours at 37 ºC to remove unligated plasmid. Following the digestion, 400 µL of EMnetik PCR cleanup beads (Beckman Coulter, C68442) were added. The sample was mixed and aliquoted into 3 x 200 µL PCR tubes. The ligated plasmid was then purified using a Beckman Coulter EMnetik according to the manufacturer's recommendations. The DNA was eluted from the beads in 100 µL of a 60% solution of Neon Transfection Buffer R and quantified using the Qubit Broad Range dsDNA kit (Invitrogen).

One million cells were electroporated for each sample using the 100 µL NEON tips. The cells were suspended in 55 µL of Buffer R. 8.56 x $10^8$ copies of ligated vector were mixed with 200 ng of integrase expression vector per million cells transfected. The volume of this mixture was adjusted to 55 µL per transfection with Buffer R. To transfect, 55 µL of the cell suspension and 55 µL of the plasmid mixture were combined. This was drawn into a 100 µL tip and the cells were electroporated with a setting of 1350 volts, 30 millisecond pulse width two times. Immediately after electroporation the cells were placed in 2 mL of complete media. After the electroporations were complete, the cell suspensions were equally divided into two wells of a 6-well plate. One well contained 10 µM ART558 (MedChem Express, HY-141520) and the other DMSO. Twenty-four hours after electroporation the media was changed. ART558 and DMSO were maintained in the media. 48-hours after transfection the cells were harvested, and genomic DNA purified using the Macherey-Nagel NucleoSpin Tissue kit (TaKaRa).

For immunofluorescence experiments with interstrand crosslink plasmids, 3.64 x $10^8$ copies of either 0, 1, or 2 ICL plasmid were transfected into hTERT-RPE1 $TP53^{-/-}$ + $PolQ^{-/-}$ + TetON Halo-Polθ cells using the Lipofectamine 3000 (Invitrogen) transfection kit with or without 200 ng of pGGL-2 integrase plasmid. Cells were treated with 15 ng/mL dox to induce Halo-Polθ expression beginning 48h prior to fixation. Cells were incubated with Jenealia Farm Halo ligand 669 for 6h prior to fixation. Fixation was completed 24h after ICL plasmid transfection. MMC-treated positive control cells were treated with 100 ng/mL MMC for 6h prior to cell fixation.

**NGS Library Preparation and Sequencing of Targeted ICL Repair Products**

100 ng of purified genomic DNA was amplified with primer pool R1V2 or R2V2 in a 100 µL reaction using the cycling conditions shown below using Phusion Polymerase (Invitrogen, M0530). Approximately half the samples were amplified with R1V2, and the remaining samples were amplified with R2V2. Following amplification, the samples were purified using the EMnetik PCR Cleanup kit with the following modification. One half of the recommended volume of beads were added to preferentially bind the genomic DNA present. The supernatant was transferred to a clean tube and the remaining volume of beads were added to preferentially bind the PCR product. The remaining steps followed the manufacturers' recommendations. The purified DNA was eluted into 50 µL of elution buffer. 20 ng of the libraries were indexed using the Illumina Dual Combinatorial Indexing protocol. The indexed PCR products were purified using the EMnetik PCR Cleanup kit according to the manufacturer's recommended protocol. The individual, purified, libraries were validated by running 2 µL of a 1:10 dilution of each on an Agilent high sensitivity D1000 TapeStation tape. The purified libraries were quantified using the Qubit Broad Range dsDNA kit (Invitrogen, Q32853) and pooled at equal molar concentrations. The pooled libraries were diluted to 70 pM in water. Twenty µL of the diluted pool was loaded onto an iSeq100 according to the manufacturer's recommendation.

**Targeted ICL Repair Sequence Analysis**

The FASTQ files were processed with ScarMapper v2.0 (*2*) to group the repair products that result in mutations and calculate the frequencies of each. The program is run by providing sgGGL-65 as a faux sgRNA guide and a k-mer length of 18. This sequence causes ScarMapper to analyze the FASTQ from a point equidistant from the two in-vitro crosslinks. The 18-nucleotide k-mer extends past the crosslink sites, allowing ScarMapper to determine if there is a substitution at the crosslink. Scars identified as non-MicroHomology deletions or Insertions were manually curated to ensure the correct identification.

**Droplet Digital PCR (ddPCR)**
ddPCR was performed as previously described except for the following changes (*14*). Forward and reverse primers were pooled together at 22.5 µM each. The working probe concentration was 6.25 µM. These changes allowed 1 µL of each to be used per reaction. In addition, the reactions using the Repaired primer/probe set were run as a duplex reaction with the genomic quantification primer/probe set. The total amount of genomic DNA in each well was 20 ng. The ddPCR plates were set up using an Opentrons OT-2 robot. Droplets were generated with the Bio-Rad QX200 AutoDG Droplet Generator. After the plate was sealed, the PCR was run according to the PCR parameters below on a Bio-Rad C1000 thermocycler. The droplets were analyzed on a Bio-Rad QX200 droplet reader with QX Manager v2.2 software.

**Imaging and analysis**
Immunofluorescence slides were imaged using Olympus BX61 microscope with HBO 100 mercury lamp and Hamamatsu ORCA-R2 camera, and 60X/1.42 Oil PlanApo N Olympus objective. Collection channels included DAPI (FF409-DI02 dichroic mirrors, FF01-377/50 excitation filter, FF02-447/60 emission filter), FITC (FF506-DI02 dichroic mirrors, FF01-482/35 excitation filter, FF01-536/40 emission filter), TxRed (FF593-DI02 dichroic mirrors, FF01-562/40 excitation filter, FF01-624/40 emission filter), and Cy5 (FF660-DI01 dichroic mirrors, FF01-628/40 excitation filter, FF01-692/40 emission filter). Fiji ImageJ was used to quantify foci. Quantification consisted of generating two duplicates of foci-containing fields and applying Gaussian blur of sigma 1 or 2. The Difference of Gaussians was generated by subtracting grayscale pixel intensity values of image 1 from image 2. The threshold was set manually for the resulting image to capture the foci while minimizing background signal. This threshold was then applied to all images, streamlined by ImageJ macro, ORION, developed in this study (https://github.com/Gaorav-Gupta-Lab/Orion.git).
Colocalization experiments were imaged using Zeiss LSM900 confocal microscope with Plan-Apochromat 63X/1.4 Oil DIC M27 objective. Images were acquired with a pinhole size set at 1AU for the longest wavelength fluorophore and kept consistent for other fluorophores, pixel size of 0.071um x 0.071um, and 541x541 pixels per image. Channels collected included DAPI (353nm excitation wavelength, 465 emission wavelength, 410-510nm detection wavelengths), Alexa488 (493nm excitation wavelength, 493nm emission wavelength, 510-595 detection wavelengths), and Alexa594 (590nm excitation wavelength, 618 emission wavelength, 595-700nm detection wavelengths). Colocalization analysis was performed using Imaris 10 software. Nuclear regions of interest (ROIs) were generated using the detect cell function. The threshold was manually set on each image to capture the DAPI signal. The spots function was used to identify RAD51 and Halo-Polθ foci in their respective channels. Spots were identified using a manually set threshold, to capture foci while minimizing background and expected spot diameter was set at 0.25 µm and 0.20 µm, respectively. The number of RAD51 foci within 0.15 µm of a Halo-Polθ foci, which we designated as colocalized, was calculated. RAD51 spot intensity was also recorded.

**Resource Table**

| Antibody | Source | Identifier |
|---|---|---|
| Rabbit Anti-ALFA | Cedarlane | N1581 |
| Mouse Anti-Halo | Promega | G9281 |
| Mouse Anti-RAD51 mab (14B4) | Novus Biologicals | NB100-148 |
| Rabbit Anti-RAD51 | Cell Signaling | 8875 |
| Goat Anti-Mouse IgG (H+L) Alexa Fluor™488 | Thermo Fisher Scientific | A-11001 |
| Goat Anti-Rabbit IgG (H+L) Alexa Fluor™488 | Thermo Fisher Scientific | A-11008 |
| Goat Anti-Mouse IgG (H+L) Alexa Fluor™594 | Thermo Fisher Scientific | A-11032 |
| Goat Anti-Rabbit IgG (H+L) Alexa Fluor™594 | Thermo Fisher Scientific | A-11012 |
| IRDye 800CW Goat Anti-Mouse | LICORbio | 926-32210 |
| IRDye 680CW Goat Anti-Rabbit | LICORbio | 926-68071 |
| Rabbit Anti-Lamin B1 (D9V6H) | Cell Signaling | 13435 |
| Mouse Anti-PCNA (PC10) | Cell Signaling | 2586T |
| Rabbit Anti-Mre11 | Cell Signaling | 4895S |
| **Software** | **Source** | |
| GraphPad Prism v9.5 | GraphPad | https://www.graphpad.com/ |
| Python version 3.10 | Python Software Foundation | https://www.python.org |
| Burrows Wheeler Aligner | (*15*) | https://github.com/lh3/bwa |
| Atropos Sequence Trimmer | (*16*) | https://github.com/jdidion/atropos |
| Völundr | (*14*) | https://github.com/Gaorav-Gupta-Lab/Volundr |
| Picard Tools | | https://broadinstitute.github.io/picard/ |
| ScarMapper | (*14*) | https://github.com/Gaorav-Gupta-Lab/ScarMapper |
| Samtools | (*17*) | http://www.htslib.org/ |
| Imaris | Oxford Instruments | https://imaris.oxinst.com/versions/10 |
| ORION ImageJ Macro | This study | https://github.com/Gaorav-Gupta-Lab/Orion |
| Brass v6.3.4 | | github.com/cancerit/BRASS |

**Supplementary Text**

### PCR Conditions for NGS Library Amplification

1. 98.0 °C for 60 seconds.
2. 98.0 °C for 15 seconds.
3. 64.0 °C for 40 seconds; -0.5 °C per cycle
4. 72.0 °C for 45 seconds.
5. Go To Step 2; 12x more.
6. 98.0 °C for 15 seconds.
7. 64.5 °C for 35 seconds.
8. 72.0 °C for 50 seconds.
9. Go To Step 6; 25x more.
10. 72.0 °C for 300 seconds.
11. 10 °C HOLD

### Illumina Dual Indexing PCR Conditions

1. 95 °C for 60 seconds.
2. 95 °C for 30 seconds.
3. 72 °C for 45 seconds.
4. Go To Step 2; 6x more.
5. 72 °C for 5 minutes.
6. 10 °C for HOLD

### ddPCR Conditions Targeted ICL Repair

1. 98 °C for 10 minutes.
2. 98 °C for 10 seconds. Ramp 3 ºC/s
3. 60 °C for 90 seconds. Ramp 3 ºC/s
4. Go To Step 2; 45x more.
5. 98 °C for 10 minutes. Ramp 3 ºC/s
6. 10 °C for HOLD. Ramp 3 ºC/s

### RT-PCR Conditions

cDNA synthesis
Template RNA 1 µg
10X dsDNase Buffer 1 µL
dsDNase 1 µL
Water to 10 µL
Incubate at 37 °C for 2 minutes
10 °C hold

5X reaction mix 4 µL
Maxima Enzyme mix 2 µL
Water 4 µL
25 °C for 10 minutes
56 °C for 15 minutes
85 °C for 5 minutes
10 °C hold

qPCR
SYBR Green master mix 5 µL
10 µM paired primers 2 µL
Water 2 µL
cDNA 1 µL

### PCR Conditions for PolQ mutants

1x TaKaRa GXL Polymerase Master Mix
500 nM primer pair
1 ng template DNA
Sterile water to desired reaction volume

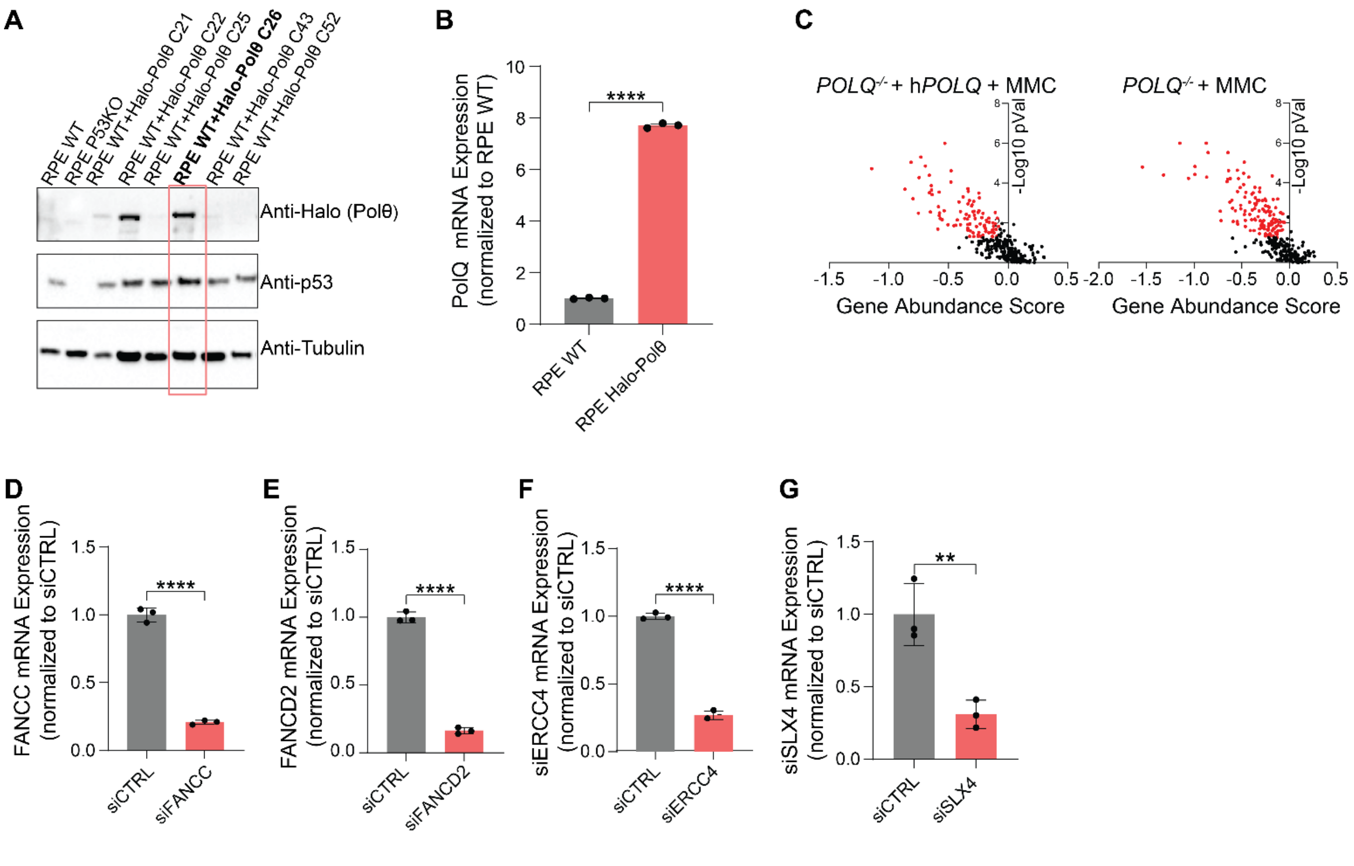

**Fig. S1.**

**Data Supporting Figure 1.**

(A) Western blot of hTERT RPE1 cells clones engineered to express Halo-tagged Polθ with anti-Halo, anti-p53, and anti-Tubulin antibodies. Clone 26 was selected for further experiments. (B) PolQ mRNA expression in hTERT RPE1 and hTERT RPE Halo-Pol θ cells determined using RT-PCR and normalized to actin (****P < 0.0001, Welch's t-test). (C) Volcano plots of gene abundance scores in hTERT RPE *TP53*$^{-/-}$ and hTERT RPE *TP53*$^{-/-}$ *POLQ*$^{-/-}$ cells after MMC treatment. (D) FANCC mRNA expression in hTERT-RPE1$^{\text{Halo-Pol}\theta}$ after treatment with non-targeting control or FANCC targeting siRNA, determined by RT-PCR and normalized to actin (****P < 0.0001, Student's t-test). (E) FANCD2 mRNA expression in hTERT-RPE1$^{\text{Halo-Pol}\theta}$ after treatment with non-targeting control or FANCD2 targeting siRNA, determined by RT-PCR and normalized to actin (****P < 0.0001, Student's t-test). (F) ERCC4 mRNA expression in hTERT-RPE1$^{\text{Halo-Pol}\theta}$ after treatment with non-targeting control or ERCC4 targeting siRNA, determined by RT-PCR and normalized to actin (****P < 0.0001, Student's t-test). (G) SLX4 expression in hTERT-RPE1$^{\text{Halo-Pol}\theta}$ after treatment with non-targeting control or SLX4 targeting siRNA, determined by RT-PCR and normalized to actin (**P < 0.01, Student's t-test).

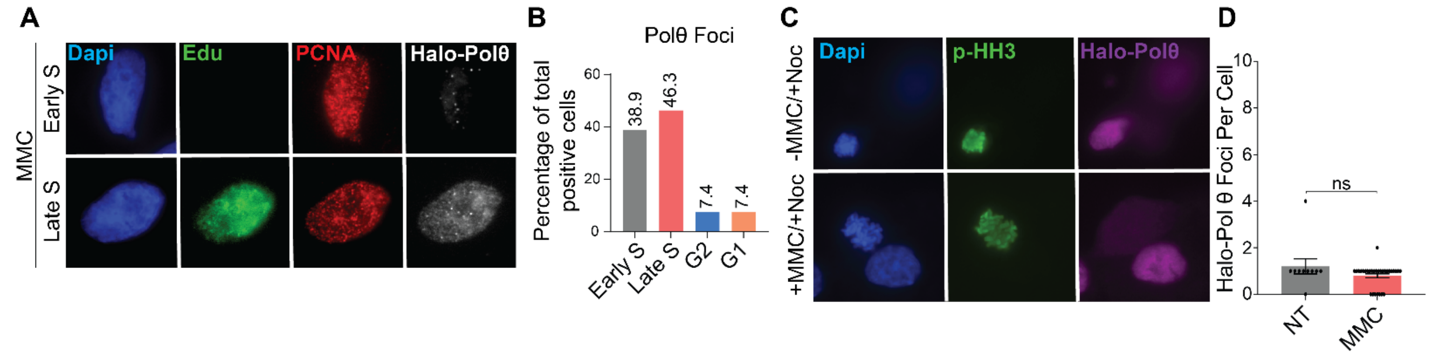

**Fig. S2.**

**Polθ Foci Form Predominately in S Phase Cells**

(A) Representative images of hTERT-RPE1$^{\text{Halo-Pol}\theta}$ cells following incubation with Edu for 30 minutes, HaloTag Ligand JF669 and 100 ng/mL MMC for 6 hours, and processing for immunofluorescence. Images show DAPI (blue), Edu (green), PCNA (red), and Halo-Polθ (grey). (B) Quantification of percentage of Polθ positive cells (determined by greater than 4 foci), in each cell cycle phase after MMC induction in hTERT-RPE1$^{\text{Halo-Pol}\theta}$ cells. (C) Representative images of hTERT-RPE1$^{\text{Halo-Pol}\theta}$ cells following incubation with HaloTag Ligand JF669, 100ng/mL Nocodazole, with or without 100ng/mL MMC for 6 hours, and processing for immunofluorescence. Images show DAPI (blue), phospho-Histone H3 (green), and Halo-Polθ (purple). (D) Quantification of Halo-Polθ foci per cell, with $\log_2$-transformed counts (n + 1) (not significant, Welch's t-test).

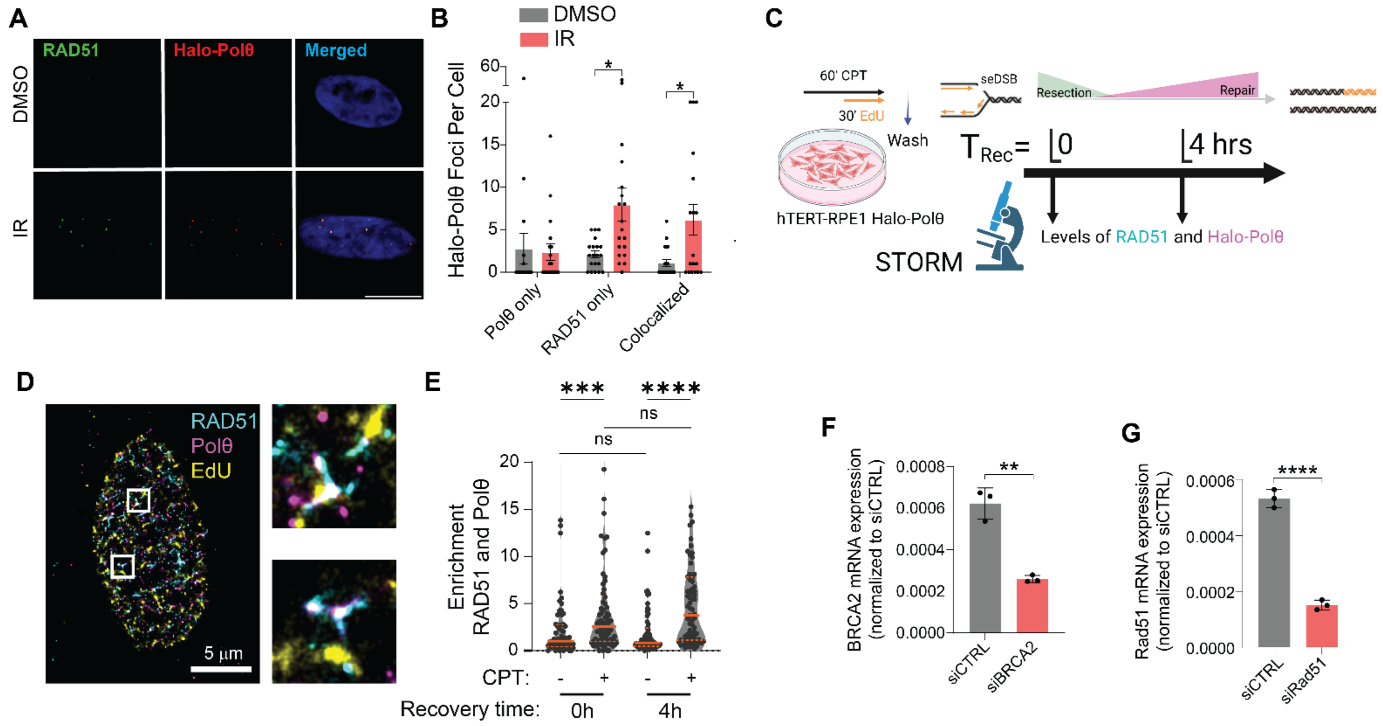

**Fig. S3.**

**PolQ colocalizes with Rad51 after different types of DNA damage**

(A) Representative images of hTERT-RPE1$^{Halo\text{-}Pol\theta}$ cells following incubation with HaloTag Ligand JFX646 and either DMSO control or 10 gray irradiation and processing for immunofluorescence. Images show DAPI (blue), RAD51 (green), and Halo-Polθ (red). (B) Quantification of Halo-Polθ foci per cell, with $\log_2$-transformed counts (n + 1) (*P <0.05, Welch's t-test). (C) Overview of STORM workflow after camptothecin (CPT) treatment. D) Representative STORM images of hTERT-RPE1$^{Halo\text{-}Pol\theta}$ cells after treatment with 100 nM CPT for 1 hour, showing RAD51 (blue), Halo-Polθ (purple), and EdU (yellow). Scale bar represents 5μm. (E) Quantification of Rad51 and Halo-Polq colocalization at 0h and 4h after CPT treatment. ***P < 0.001, ****P < 0.0001, two-tailed Mann-Whitney test. (F) BRCA2 mRNA expression in hTERT-RPE1$^{Halo\text{-}Pol\theta}$ after treatment with non-targeting control or BRCA2 targeting siRNA, determined by RT-PCR and normalized to siCTRL (**P < 0.01, two-tailed Student's t-test). (G) RAD51 mRNA expression in hTERT-RPE1$^{Halo\text{-}Pol\theta}$ after treatment with non-targeting control or RAD51 targeting siRNA, determined by RT-PCR and normalized to siCTRL (**P < 0.01, two-tailed Student's t-test).

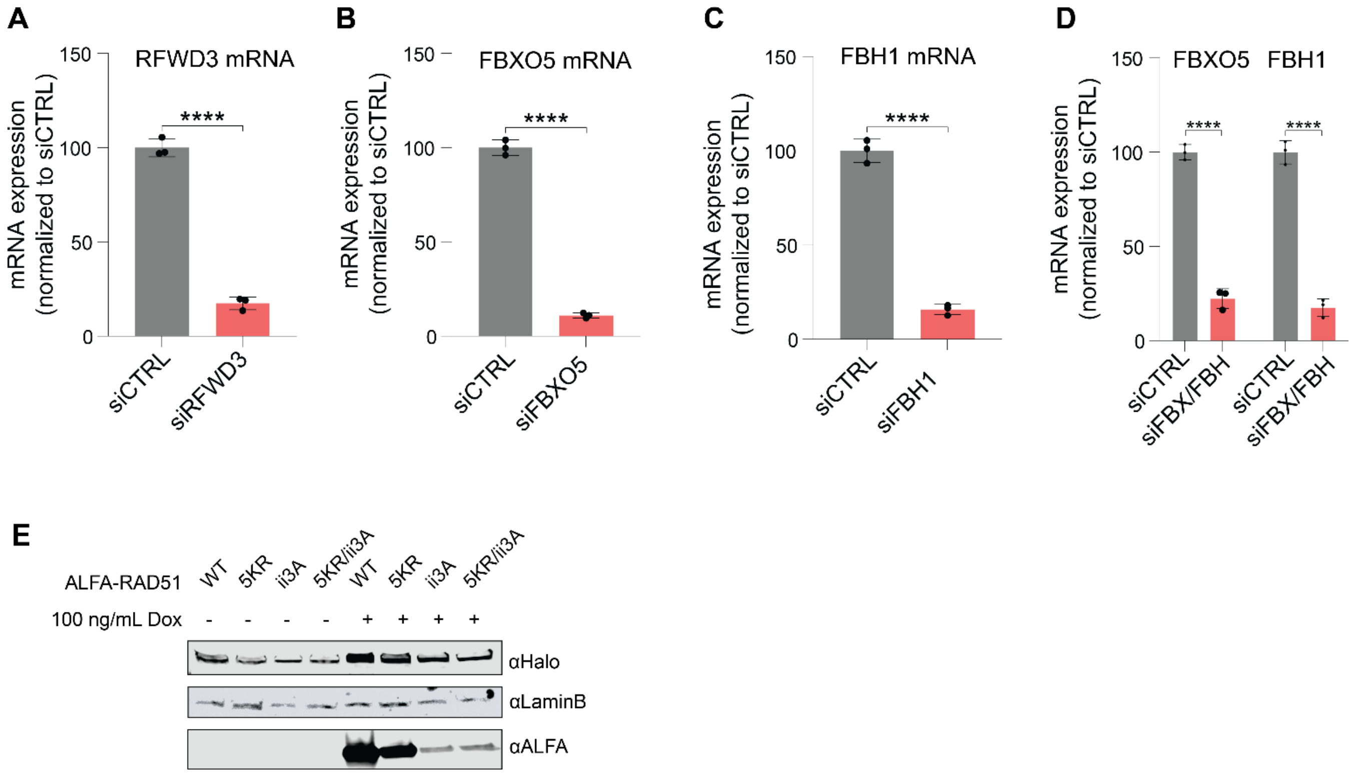

**Fig. S4.**

**Ubiquitylation is Required for Interaction with Polθ**

(A) RFWD3 mRNA expression in hTERT-RPE1$^{Halo\text{-}Pol\theta}$ cells after knockdown of non-targeting control (siCTRL) or RFWD3 (siRFWD3), determined using RT-PCR and normalized to actin (****P < 0.0001, Student's t-test). (B) FBXO5 mRNA expression in hTERT-RPE1$^{Halo\text{-}Pol\theta}$ cells after knockdown of non-targeting control (siCTRL) or FBXO5 (siFBXO5), determined using RT-PCR and normalized to actin (****P < 0.0001, Student's t-test). (C) FBH1 mRNA expression in hTERT-RPE1$^{Halo\text{-}Pol\theta}$ cells after knockdown of non-targeting control (siCTRL) or FBH1 (siFBH1), determined using RT-PCR and normalized to actin (****P < 0.0001, Student's t-test). (D) FBXO5 and FBH1 mRNA expression in hTERT RPE Halo-Polθ cells after knockdown of non-targeting control (siCTRL) or both FBXO5 and FBH1 (siFBX/FBH), determined using RT-PCR and normalized to actin (****P < 0.0001, Student's t-test). (E) Chromatin extracts of hTERT-RPE1$^{Halo\text{-}Pol\theta}$ cells showing protein expression of Doxycycline-inducible ALFA-RAD51 mutant constructs.

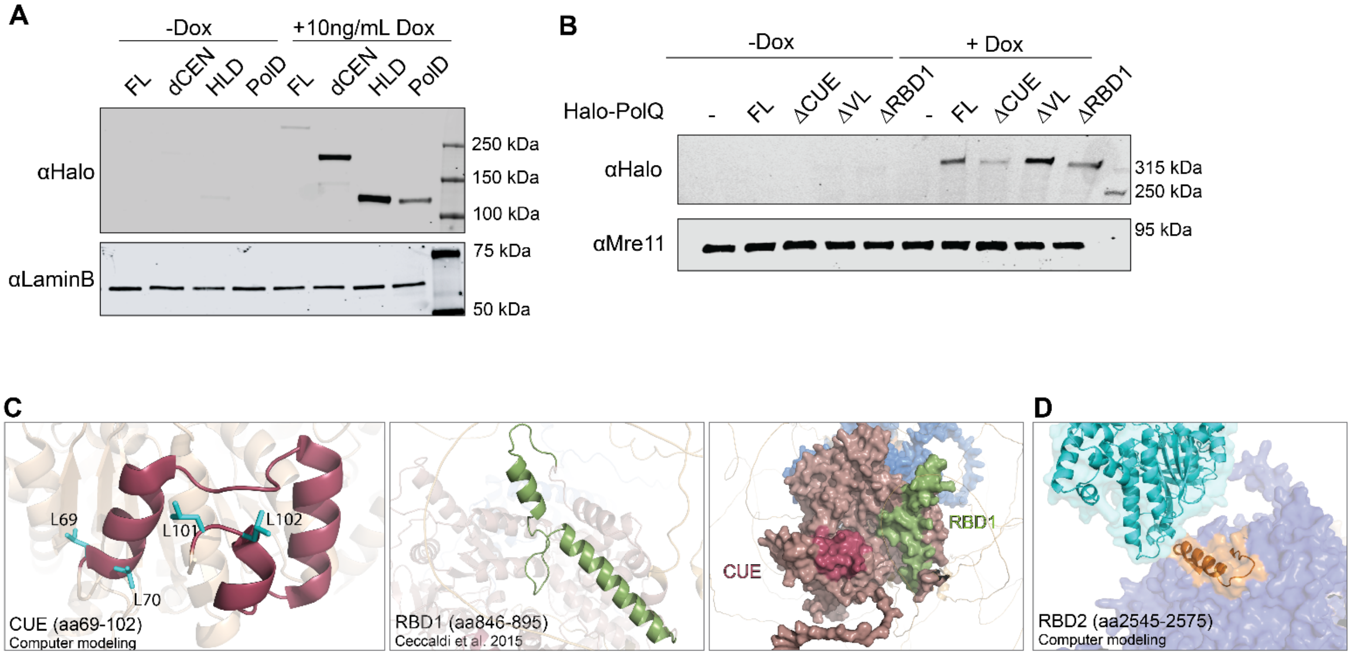

**Fig. S5.**

**Polθ RAD51 and ubiquitin binding-domains are required for interaction**

(A) Nuclear extracts of hTERT-RPE1$^{Halo\text{-}Pol\theta}$ cells showing protein expression of Doxycycline-inducible Halo-Polθ mutant constructs, full length protein (FL), central domain deletion (ΔCEN), helicase domain only (HLD), and polymerase domain only (PolD). (B) Nuclear extracts of hTERT-RPE1$^{Halo\text{-}Pol\theta}$ cells showing protein expression of Doxycycline-inducible Halo-Polθ mutant constructs, full length (FL), CUE domain deletion (Δ CUE), dimerization domain deletion (ΔVL), and RAD51 binding domain deletion (Δ RBD1). (C)PyMOL3 cartoon representation of CUE domain (left) with Leucines mutated in 4LA highlighted in cyan, RBD1 (middle), and structural location of both domains in surface representation (right). (D) Top HADDOCK v2.4 cluster for interaction between full length Halo-Polθ and RAD51 shows interaction at RBD2. HADDOCK score of -70 represents high confidence of interaction.

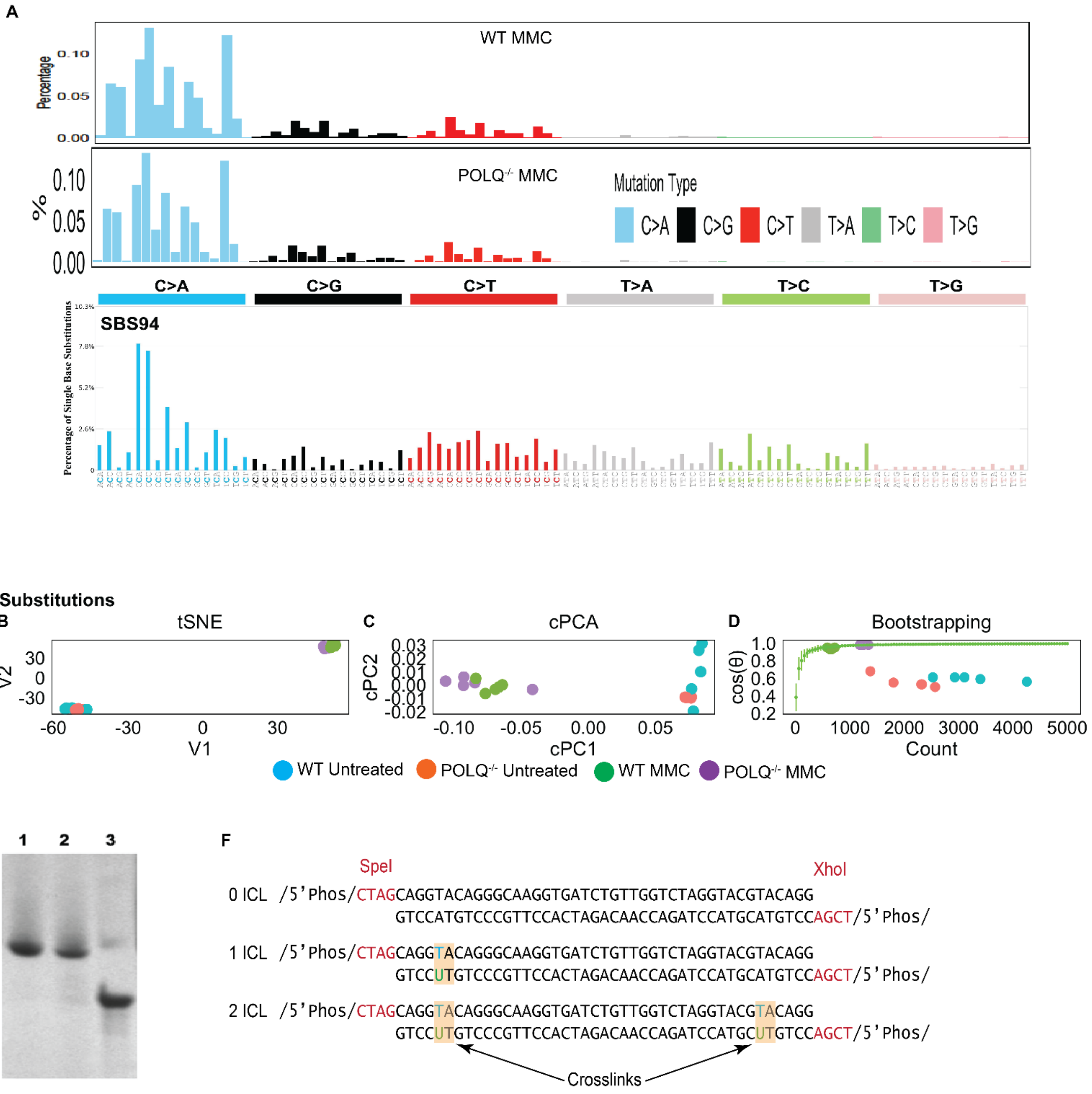

**Fig. S6.**

**Data Supporting Figure 5.**
(A) tSNE plot showing that the SNV spectrum in untreated and Mitomycin C treated in hTERT RPE *TP53*$^{-/-}$ cells (WT) and hTERT RPE *TP53*$^{-/-}$ *POLQ*$^{-/-}$ cells (*POLQ*$^{-/-}$ ) is not dependent on Polθ expression. (B) cPCA plot showing that the SNV spectrum in untreated and Mitomycin C treated in hTERT RPE *TP53*$^{-/-}$ cells (WT) and hTERT RPE *TP53*$^{-/-}$ *POLQ*$^{-/-}$ cells (*POLQ*$^{-/-}$ ) is not dependent on Polθ expression. (C) Bootstrap analysis plot showing that the SNV spectrum in untreated and Mitomycin C treated in hTERT RPE *TP53*$^{-/-}$ cells (WT) and hTERT RPE *TP53*$^{-/-}$ *POLQ*$^{-/-}$ cells (*POLQ*$^{-/-}$) is not dependent on Polθ expression. (D) COSMIC plots showing that Mitomycin C treated in hTERT

RPE *TP53*$^{-/-}$ cells (WT MMC) and hTERT RPE *TP53*$^{-/-}$ *POLQ*$^{-/-}$ cells (*POLQ*$^{-/-}$ MMC) have the same mutation spectrum and that this matches COSMIC SBS94, the Mitomycin C SNV spectrum in human cells. (E) 8% denaturing PAGE Urea gel: Lane 1: 1-ICL; Lane 2: 2-ICL; Lane 3: control single strand oligonucleotide. (F) Sequences of oligonucleotides with click chemistry cross links.
